# Multi-immersion Oblique Plane Microscope (miOPM): A reconfigurable platform for high-resolution Light-Sheet Fluorescence Microscopy

**DOI:** 10.1101/2025.10.04.680473

**Authors:** Bingying Chen, Alfred Millett-Sikking, Seweryn Gałecki, Stephan Daetwyler, Jenny Jiou, Jim Monistrol, Qionghua Shen, Felix Zhou, Hsin-Yu Lin, Edward Jenkins, Miriam C. Stein, Madeleine Marlar-Pavey, Gabriel Sturm, Alexander L. Li, Qiaosi Tang, Brian Feng, Ulises Diaz, Yannan Chen, Ary Shalizi, Astrid Gillich, Jonathan R. Friedman, Raju Tomer, Bo-Jui Chang, Wallace F. Marshall, Sarah Shahmoradian, Kevin M. Dean, Reto Fiolka

## Abstract

Light-Sheet Fluorescence Microscopy (LSFM) enables gentle, rapid, and efficient volumetric imaging of biological specimens. Despite its potential, LSFM has fragmented into multiple variants, each narrowly optimized for specific samples or imaging regimes, limiting its broad applicability. To overcome this barrier, we introduce the multi-immersion oblique plane microscope (miOPM), an adaptable LSFM platform for high-resolution imaging ranging from subcellular dynamics to whole organisms and cleared tissues. miOPM uniquely supports seamless interchangeability of oil, water, and air objectives, and maintains diffraction limited performance across a refractive index range of 1.33 - 1.51. It is compatible with standard sample mounting, current clearing protocols and high-throughput 3D imaging. We leverage miOPM to image diverse biological specimens at multiple spatial scales. By greatly expanding the adaptability and ease of use of LSFM, miOPM is poised to democratize advanced three-dimensional imaging.

## Main

Light-Sheet Fluorescence Microscopy (LSFM) has established itself as a major fluorescence microscopy technique due to its gentle, high-speed volumetric imaging capabilities^1,2^. It enables imaging of subcellular dynamics, intravital imaging in small model organisms, and *in toto* imaging of cleared mouse and human tissues^3–5^. However, a limiting factor in the widespread adoption of LSFM is its lack of adaptability and versatility. Often, specialized sample mounting is required, and each light-sheet instrument is optimized for a narrow application space. For example, high-resolution variants of LSFM, such as Lattice Light-Sheet Microscopy (LLSM)^6^ and Field Synthesis^7^, use a high NA detection objective paired with a custom illumination objective designed specifically to fit into the remaining mechanical space. Both objectives also require long working distances to ensure the overlap of the light sheet and detection focal plane (**Figure 1A**). The orthogonal objective geometry inherently limits the maximum solid angle for collecting emitted light, and the objectives cannot readily be swapped to reconfigure the microscope for a different application. Consequently, while high-resolution LSFM microscopes have been widely adopted for studying subcellular dynamics in single cells, they are not well suited for whole-embryo imaging − a need met by an entirely different class of LSFM systems^2,8^. Likewise, imaging cleared tissues^9^ demands specialized LSFM configurations, sometimes even tailored to a particular clearing technique^10^.

**Figure 1:**
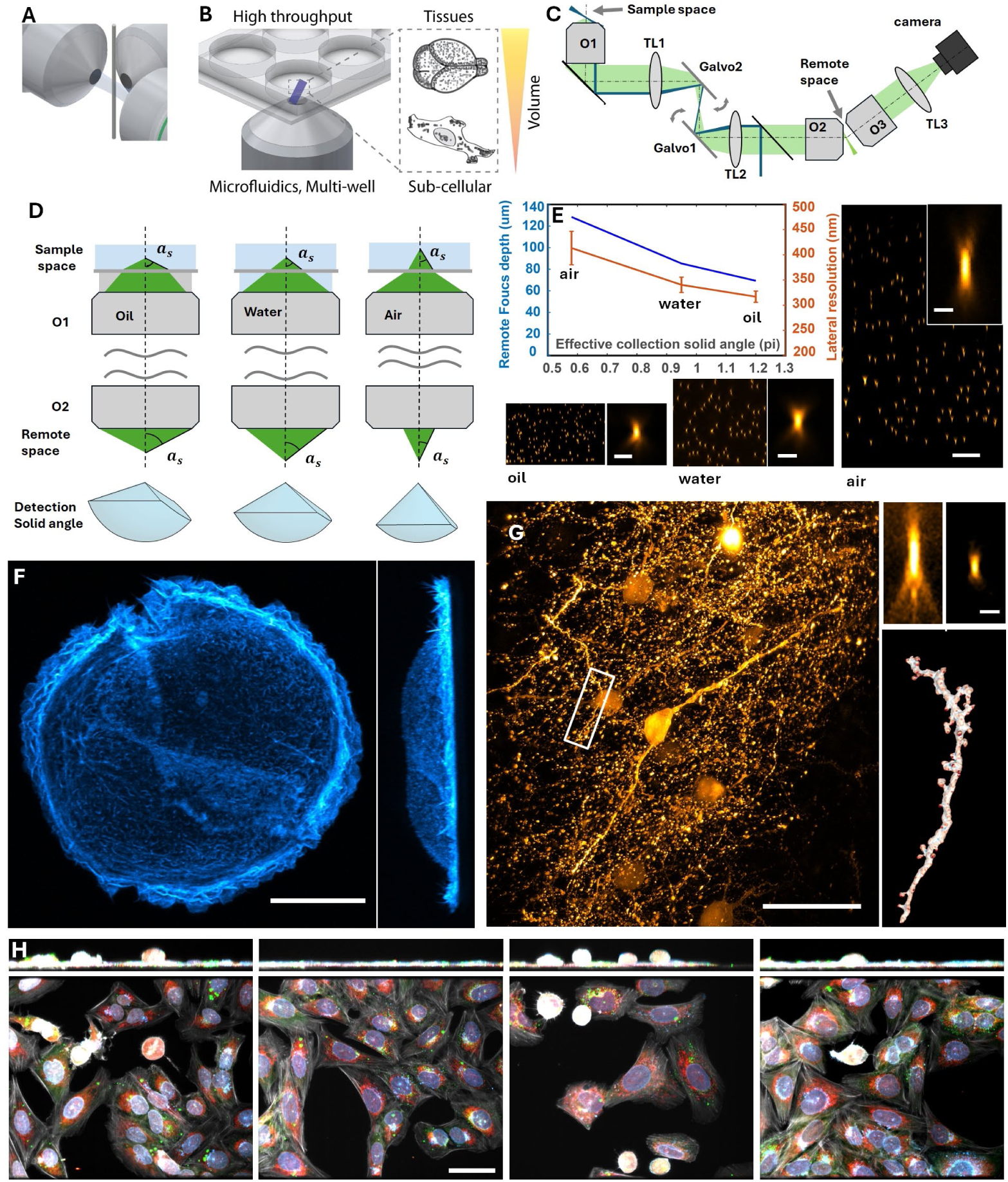
Light-sheet fluorescence microscopy (LSFM) architectures. **A** Orthogonal arrangement of illumination and detection objectives in conventional LSFM. **B** Multi-well and multi-scale imaging with multi-immersion oblique plane microscopy (miOPM). **C** Optical layout of miOPM. Blue depicts the excitation light-sheet, green the detection path with the primary (O1), secondary (O2) and tertiary (O3) objectives and the tube lenses (TL1, TL2, TL3). **D** Remote focusing in the presence of refractive index mismatch. Diffraction limited performance can be maintained if angles (αs) in sample space are mapped to remote space identically. Relay optics between the primary (O1) and secondary objective (O2) are left out for brevity. **E** Remote focusing range and lateral resolution as a function of the collected solid angle. Lateral resolution estimated from imaging fluorescent nanospheres in agarose (n>280). Insets show magnified point spread functions. **F** Maximum intensity projections (x-y, left; x-z right) of a SU8686 cell labeled with Tractin-mRuby as imaged by OPM with an oil immersion objective. **G** Maximum intensity projection of clarity cleared mouse brain section, labeled for Thy1-eGFP neurons, as imaged by OPM with a silicone oil objective. Insets: top: PSF before and after optimizing miOPM to Clarity media. Bottom: segmented axon from the boxed area in **G**. **H** Contact-less multi-well imaging with an air objective. Four maximum intensity projections (x-y, bottom; y-z, top) of selected volumes out of 384 wells that were visited, containing fixed U-2 OS cells stained with Mitotracker Far Red, Hoechst, Phalloidin CF430, Phenovue 493, and WGA-Alexa555.Scale bar: **E** 1 & 10 μm; **F** 20 μm; **G & H** 50 μm; **G** inset 1 μm.

In contrast to traditional LSFM designs, oblique plane microscopy (OPM) uses a single objective for both illumination and detection^11^, eliminating the need to carefully match the illumination and detection objectives (**Figure 1B**). However, this apparent flexibility comes with its own constraints: the downstream optics, which are necessary to map the tilted light-sheet image onto a camera in a distortion-free and diffraction-limited manner, need to be matched to the primary objective^12^. Consequently, changing the primary objective in OPM previously required changing the entire downstream optical train (**Figure 1C**), which in practice means building a new microscope.

As a result, laboratories must often acquire and operate multiple LSFM instruments, many of which require dedicated expertise to use effectively. Even in advanced microscopy facilities, it is challenging to maintain the breadth of LSFM systems necessary to enable widespread biological imaging applications. Thus, users must revert to other, potentially sub-optimal imaging techniques or change their experimental design.

To address these challenges, we introduce the multi-immersion oblique plane microscope (miOPM), a single LSFM system that accommodates applications ranging from imaging sensitive sub-cellular dynamics with high numerical aperture (NA) oil objectives to imaging larger specimens with long-working-distance water or air objectives, such as model organisms or cleared tissue sections. Building upon the Any Immersion Remote Focusing (AIRR) framework^13^, which we formally derive in this manuscript (**Supplementary Note 1**), miOPM enables objectives with the same effective focal length (EFL) to be freely interchanged, regardless of immersion media (**Figure 1D-H**). We demonstrate this principle using Nikon’s 40x objective family (EFL = 5 mm), spanning oil, silicone, water, and air immersions with diverse NAs and working distances, thereby allowing resolution, sensitivity, and axial coverage to be tailored to experimental needs (**Figure 1E-H**, **Table 1**, **Supplementary Figure 1-2**). Leveraging this advance, miOPM thus supports standard sample mounting, rapid volumetric acquisition, environmental control, contact-free high-throughput imaging in multi-well plates, and compatibility with advanced modalities such as correlative light and electron microscopy. Together, these features establish miOPM as a versatile and broadly adaptable system that consolidates previously fragmented LSFM applications within a single microscope.

**Table 1.** Properties and performance of miOPM with different primary objectives. The solid angle was computed from the theoretical light transmission, shown in **Figure 1E**. The effective NA was computed as a half angle from a cone with the solid angle in the first column. The resolution values were measured as Full width half maximum (FWHM) of 100nm fluorescent nanospheres embedded in Agarose. No deconvolution was applied, only de-skewing and rotation of the raw data. The mean and standard deviation values with each objective were calculated from 304, 328, 279, and 280 measurements within a 20-µm thick volume around the focal plane respectively. See also **Supplementary Figure 1** for spatial resolution as a function of depth. Imaging depth is based on AIRR theory. If sample R.I. matches the R.I. of any immersion objective, the remote focus range doubles.

| Objective | Collection Solid Angle | Effective NA | Imaging depth ($\mu\text{m}$) | Resolution - X (nm) | Resolution - Y (nm) | Resolution - Z (nm) |
| --- | --- | --- | --- | --- | --- | --- |
| **Air** | 0.58 pi | 0.94 | 128.4 | 448.8 ± 24.2 | 413.5 ± 33.3 | 1684.3 ± 58.6 |
| **Water** | 0.95 pi | 1.13 | 85.3 | 370.2 ± 17.9 | 340.4 ± 15.2 | 886.7 ± 63.1 |
| **Silicon oil** | 1.17 pi | 1.21 | 70.1 | 365.7 ± 19.4 | 316.6 ± 12.4 | 762.1 ± 45.0 |
| **Oil** | 1.20 pi | 1.22 | 69.3 | 358.1 ± 12.5 | 316.9 ± 11.2 | 753.6 ± 40.9 |

## Results

### Foundations of Any Immersion Remote Refocusing (AIRR)

To motivate the AIRR theory, we first revisit the principles of remote focusing: A primary and a secondary objective (O1 and O2; see also **Extended Figure 1A-B**) form a remote focusing system that produces a distortion-free and diffraction-limited 3D image of the sample. An emitter residing on the nominal focal plane of O1 (black dot) produces a flat wavefront in infinity space, which is then re-focused onto the nominal focal plane of O2. An emitter that is located by a distance z_1_ outside of the nominal focal plane (red dot) will result in a curved wavefront. When entering the secondary objective, the curvature has the opposite sign, and as such the wavefront will be re-focused at -z_2_ (in the opposite direction) from the focal plane of O2. Aberrations for this refocusing operation are minimized if the pupil magnification is equal to the ratio of refractive index surrounding the original point and the reimaged point (e.g. if a watery object is imaged into air the pupil magnification must be 0.75). With a more in-depth description in **Supplementary Note 1**, this results in diffraction limited 3D imaging over a finite depth range, and the axial and lateral magnification are identical. It also results into mapping angles in sample space identically to the remote space, i.e. if the light-sheet is tilted by 45 degrees in sample space, its image is tilted by the same amount in remote space.

**Extended Figure 1C-D** illustrates the working principle of AIRR, where there is a refractive index mismatch between the immersion media of the primary objective, and the sample space. When the interface of the refractive index boundary coincides with the nominal focal plane, an emitter in the focal plane (black dot) leads again to a flat wavefront in the infinity space, and correspondingly to a diffraction limited focus in the nominal focal plane of O2. A point to the left (outside of the nominal focal plane, deeper in the sample medium) will incur a curved wavefront. When we analytically derived the resulting wavefront error (compared to a flat wavefront), the resulting terms can largely be compensated by the conjugate wavefront distortion introduced by the secondary objective (full derivation of the terms in **Supplementary Note 1**). The result of our theoretical modeling of AIRR is that for the best balancing of aberrations, one needs to fulfil a similar magnification criterion as in conventional remote focusing: the lateral magnification needs to equal the ratio of the sample’s refractive index to the index of refraction of the secondary objective. Like in traditional remote focusing, angles in sample space are also mapped to the same angle in remote space, as shown in **Figure 1D**.

Consequently, the refractive index of the immersion objective does not matter in the AIRR framework. Therefore, objectives with any immersion medium can be used interchangeably, as long as they provide the same magnification to the remote focus system and possess a sufficient angular aperture to launch a light-sheet at the desired oblique angle in sample space. Additionally, if the immersion and sample refractive index differ, then the interface (typically the coverslip) needs to coincide with the focal plane of the primary objective. This reduces the imaging depth in half compared to traditional remote focusing (see also **Supplementary Note 1**).

### Continuous adaptation to the sample’s refractive index

A critical requirement for distortion-free remote focusing across samples of varying refractive index is the ability to adjust magnification between the sample and remote space. This can be done by engineering custom tube lenses (i.e. TL1 and TL2 in **Figure 1C**), each optimized for imaging in a specific medium. Thus, changing the refractive index of the sample requires different tube lenses, which in turn requires rebuilding and realigning the entire OPM system. To overcome this, we introduce a zoom lens that can be used in an OPM system to continuously change the magnification to the remote space (**Figure 1C and Extended Figure 2**). Importantly, the zoom lens keeps both its front and back focal planes stationary to keep the relative alignment of OPM’s optical train intact (see also **Extended Figure 2**) *and* maintains telecentric imaging. The zoom lens consists of three lens groups that move relative to each other (**Extended Figure 2 and Supplementary Figure 3**) to cover a magnification range of 1.33 to 1.51 in our miOPM setup. When the net refractive index of the sample is known, the zoom lens is adjusted to yield the corresponding magnification into the remote space. This ensures that diffraction limited imaging can be achieved (see also insets **Figure 1G**), and importantly, it also avoids distortions of the 3D volume that is acquired. This aspect is often overlooked in microscopy, i.e. when a 3D stack is acquired with a mismatched immersion objective the resulting 3D volume is non-linearly distorted in the axial dimension^14,15^. The zoom lens circumvents such distortions optically and makes miOPM compatible with common tissue clearing protocols.

**Figure 2.**
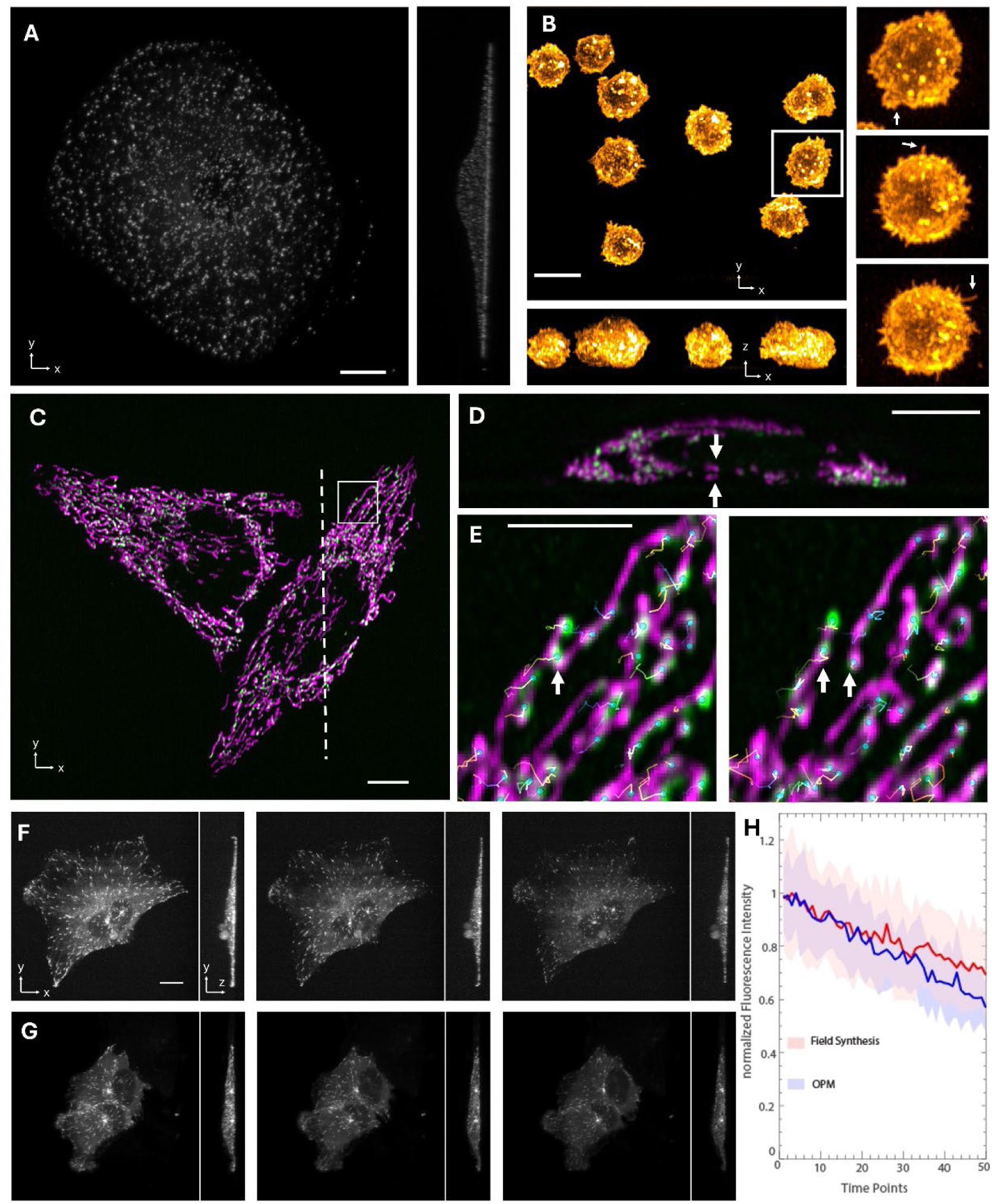
Sensitive high-resolution imaging with the oil immersion objective. **A** Clathrin coated pits, labeled with apha-eGFP adaptin, in a fixed U-2 OS cell, as imaged with the miOPM. **B** CD8+ T-cells, membrane labelled with CellMask Orange, as imaged with the miOPM. Three selected timepoints on the square boxed region shown on the right. Arrows point to protrusions and microvili **C** Mitochondria (magenta) and mitochondrial DNA (green) as imaged in a HeLa cell. **D** Axial cross section along the dotted line **C**. Arrows point at two closely space mitochondria. **E** Two selected timepoints (12s interval) on the square boxed region in **C**. Arrows point at fission event of mitochondrial DNA. **F** Three time points (first, 25^th^ and 50^th^) of a timelapse of imaging an RPE cell labeled with mNeon green using miOPM. **G** Three time points imaged with comparable acquisition parameters of the same cell line using a Field Synthesis microscope. **H** Comparison of bleaching for miOPM and Field Synthesis. Scale bars: **A,B,C,D**,**F** 10 μm; **E** 5 μm.

The zoom lens and the AIRR theory are used in this manuscript synergistically. The zoom lens is used to adapt miOPM to the refractive index of the sample (see also **Supplementary Table 1** for acquisition settings used in this manuscript). When the magnification of the zoom lens matches the refractive index of the immersion media of the objective, traditional remote focusing applies. If there is mismatch between the sample’s refractive index and the immersion media, the AIRR theory applies. A practical consequence is that when one can operate in the RF regime, there is no stringent requirement on placing the coverslip. This can be of importance for large samples that require mechanical tiling in the z-direction. In this manuscript, we highlight applications that benefit from miOPM’s ability to operate in either regime.

### Sensitive, high-resolution imaging with an oil objective

With an oil-immersion objective (40X/NA1.3), miOPM achieves its most sensitive and highest-resolution mode for imaging subcellular dynamics. In **Figure 2A**, clathrin coated pits, labeled with alpha-eGFP adaptin, in an Adult Retinal Pigment Epithelial (ARPE) cell are shown after deconvolution. As the clathrin coated vesicles form small spheres below 100nm, they allowed us to estimate the resolving power of miOPM in a cellular context. The spatial resolution, as estimated by image decorrelation, amounted to 254±17 nm throughout the stack (77 slices analyzed).

Next, we used the sensitive imaging capacity of miOPM with the oil immersion objective to capture morphological features of T-cells. T-cell protrusions, referred to as microvilli, are labile, actin-rich, 100-800nm long, ∼300-400nm diameter structures that are used to rapidly scan the membranes of target cells for antigens. The dynamics, height and diameter of these protrusions are key to T cell function. Microvillar dynamics determines antigen scanning speed, the height helps T cells overcome steric barrier function of the ‘sugar barrier’ (i.e., glycocalyx) surrounding most cells, and the diameters impacts T-cell signaling decisions^16,17^. Quantifying these features can therefore provide valuable insight into T-cell function. However, as these structures can form and collapse in the order of seconds, and given their size, high spatiotemporal imaging with low bleaching and phototoxicity is required to capture unperturbed protrusion activity. We labelled the membrane of CD8+ T-cells with CellMask Orange and volumetrically imaged them with 1s volume rate over 300 timepoints, which allowed us to observe the initial formation (arrows in **Figure 2B**) and dynamics of microvilli (**Supplementary Video 3**). Our data highlights the dynamic activity of protrusive structures, with speeds and spatial resolution sufficient for capturing their formation and movement over the T-cell surface.

To highlight the potential to image subcellular dynamics, we labeled HeLa cells for mitochondria (MitoTracker Red) and mitochondrial DNA (mtDNA, labeled with PicoGreen). mtDNA can be visualized as small puncta, which are well resolved with miOPM in 3D (**Figure 2C-D**). Using the uTrack3D tracking software^18^, we quantitatively observed translocation of mtDNA along mitochondria, and apparent splitting of punctae in the event of mitochondria fission (**Figure 2E and supplementary Video 4-5**).

Lastly, to compare the sensitivity to a conventional LSFM instrument, we imaged RPE cells labeled with EB3-mNeon green with miOPM and our previously published Field Synthesis system^7^ (**Figure 2F-G and Supplementary Video 6**), which features a traditional LSFM architecture with a NA 1.1 detection objective and generates a high duty cycle square lattice light-sheet. By comparing the 30 brightest spots in each timeframe, we established a bleaching curve for each system, which overall is comparable (**Figure 2H**).

### Water objective for imaging extended samples

Operating miOPM with a water objective (40X/NA1.15) as the primary lens extends the volumetric reach while still providing sub 400nm lateral resolution (**Table 1**). For watery samples, the water immersion objective unlocks the full working distance of the primary objective. We used this to image a four-time expanded liver slice, labeled for chromatin nucleic acid with SYTOX Green, over a height of ∼600 μm by tiling eleven volumes in the vertical direction (**Figure 3A**). In contrast to lattice light-sheet microscopy, which interfaces a slab of expanded gel at an angle^19^, here the interface to the gel is oriented normal to the optical axis of the primary objective. As such, we expect less aberrations, and miOPM affords for high-resolution imaging throughout the volume. In **Figure 3A**, one can discern in the axial cross section prominent nucleoli (non-deconvolved data is shown). We used u-Segment3D^20^ to segment individual 3D nuclei, and segmented nucleoli within each nuclei by intensity thresholding and connected component analysis. The segmented nuclei (transparent gray) and nucleoli (colored) were 3D rendered using ChimeraX (**Figure 3B and Supplementary Video 7)**.

**Figure 3.**
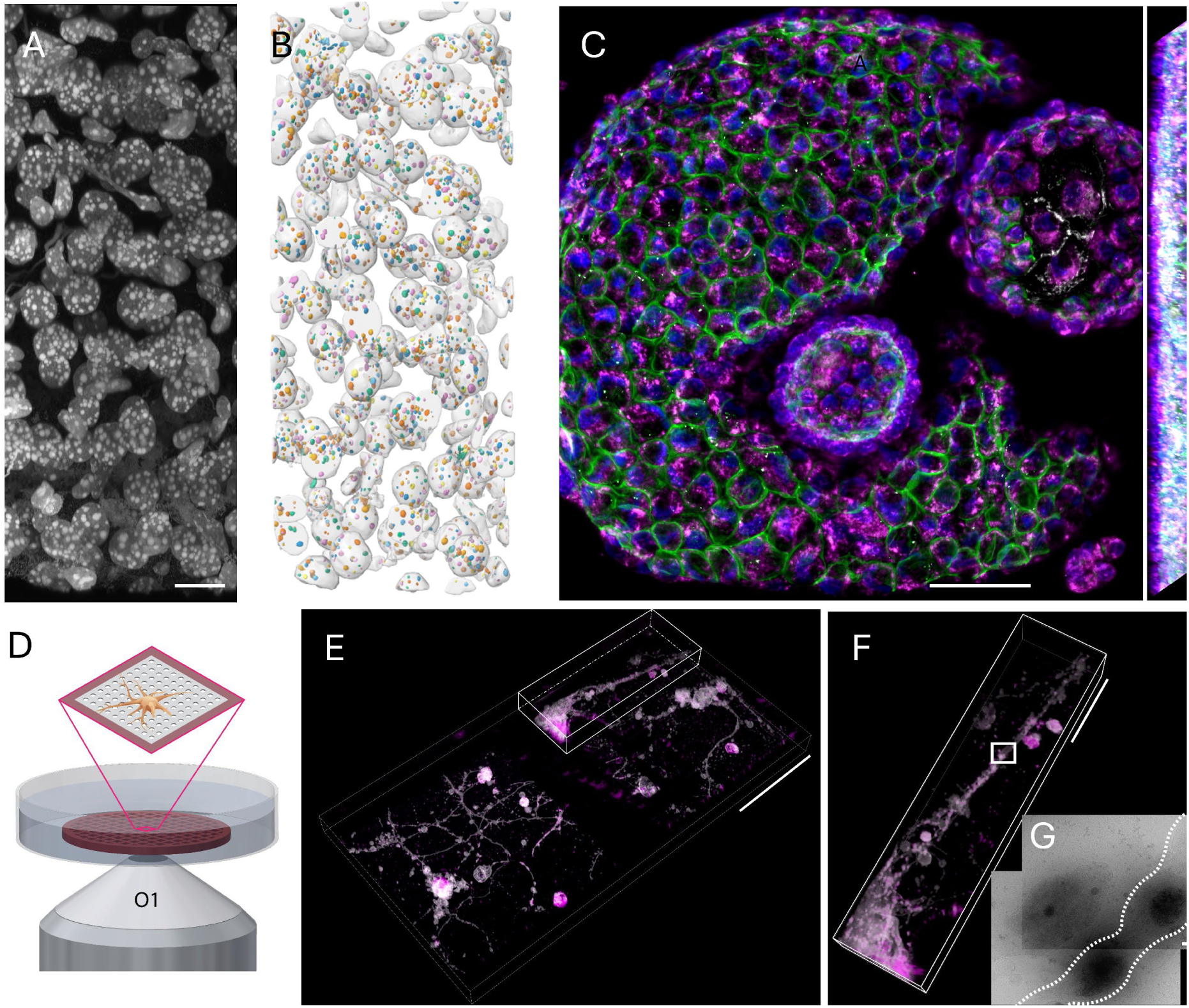
Imaging of multicellular samples and correlative imaging. **A** Nuclei, labeled with sytoxGreen, in an expanded liver tissue slice, as imaged with miOPM with a water immersion objective and using vertical tiling. Maximum intensity projection of raw (non-deconvolved) data. **B** Segmentation of the nuclei (gray) and nucleoli (color) in the volume shown in A. **C** Mouse alveolar organoids immunostained with Muc1 labeling AT2 cells (green), TOM20 labeling mitochondria (magenta), RAGE labeling AT1 cells (gray) and DAPI (blue). **D** Imaging neurons cultured on an electron microscopy (EM) grid with holey carbon support. **E** Striatal neurons atop an EM grid, labeled with Memglow488 (grey) and fluorescently-tagged alpha-synuclein protein (magenta) as imaged by miOPM. **F** Magnified view of the boxed region in **E**. **G** High-resolution transmission cryo-EM micrograph of the boxed region in **F**, white dotted lines outline the axon. Scale Bars: **A, C, E** 50 μm, **F** 20 μm, **F** 500 nm.

To illustrate the application to multicellular live samples, we imaged an organoid made from Alveolar epithelial type II (AT2) cells which were isolated from the lungs of adult C57BL6/J mice. AT2-derived organoids are a model for the function of the adult progenitor cell population in lung alveoli, and are used to study lung repair and lung disease^21,22^. To visualize mitochondrial dynamics in AT2 cells, the progenitor cells of alveoli, and their progeny (AT1 cells), the organoid was immunostained for Muc1 labeling AT2 cells, TOM20 labeling mitochondria, RAGE labeling AT1 cells and DAPI, and imaged by miOPM as shown in **Figure 3C**.

Lastly, to demonstrate the application of correlative light and electron microscopy (CLEM) for capturing dynamic molecular events of fluorescently tagged proteins—in this case, pre-formed α-synuclein fibril (PFF) “seeds”—we cultured primary mouse striatal neurons at 14 days in vitro (DIV14) directly on holey carbon–coated gold electron microscopy (EM) grids (schematic shown in **Figure 3D**). These pathological fibril “seeds” are capable of templating the misfolding and aggregation of endogenous α-synuclein, thereby modeling early events of synucleinopathy.^23,24^ To delineate the encompassing overall neuronal architecture, including fine axonal and dendritic extensions, we labeled the plasma membrane with MemGlow-488^25^. Fluorescently tagged α-synuclein PFFs (aSyn-647) were subsequently added to the culture medium to track their interaction with and uptake by neurons. While the EM grid introduced an axial offset relative to the coverslip, the use of the water-immersion objective allowed flexible axial positioning and maintained high optical sectioning strength. To explore the sample, we simultaneously imaged two adjacent grid squares (**Figure 3E**). Fluorescence imaging revealed multiple discrete α-synuclein puncta along axons, consistent with internalized fibril clusters or sites of membrane engagement (**Figure 3F** and **Supplementary Figure 4**). Immediately afterward, the grids were blotted and plunge-frozen in liquid ethane to preserve ultrastructure in vitreous ice, following established workflows for preparing cultured neurons for cryo-EM. Grids were initially screened on a 200-kV Talos Arctica, and high-resolution cryo-electron microscopy was subsequently performed on a Titan Krios (300 kV) equipped with a direct electron detector. The correlative EM imaging in **Figure 3G** and **Supplementary Figure 4** shows that the fluorescent aSyn-647 puncta correspond to fibrillar aggregates localized within neurites.

### Air objective for contact-less large volume imaging

The use of an air objective enables miOPM to perform contact-less, yet distortion free volumetric imaging. The use of the air objective also results in the largest optically scanned (i.e. without mechanical tiling) volume owing to its large remote focusing height (**Table 1**). When used with a short light-sheet, the air objective allows miOPM to still distinguish between organelles, such as mitochondria in 3D (**Figure 4A**, non-deconvolved data shown).

**Figure 4.**
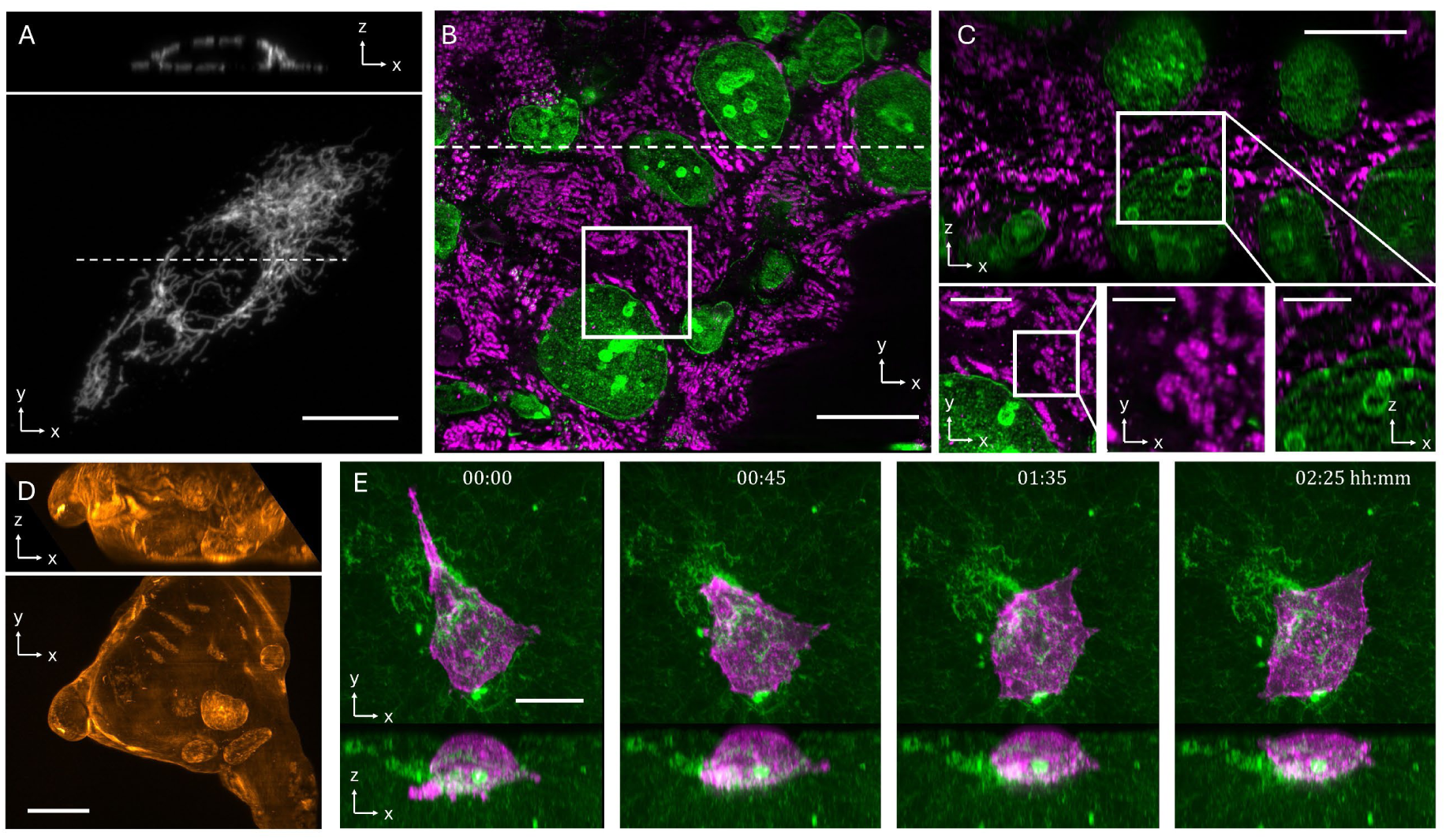
Contact-less 3D imaging of biological samples with an air primary objective. **A** U-2 OS cell labeled with EGFP-OMP25. An axial cross section along the dotted line is shown above. **B** Expanded hepatocellular carcinoma specimen stained for mitochondria (Hsp60, magenta) and nuclei (SYTOX Green and Lamin A/C, green). **C** Cross-sectional view along the dotted line in B. Insets show magnified versions of lateral and cross-sectional views. **D** Amoeba C*haos carolinensis* labeled with lifeact-GFP2. **E** Imaging SU8686 cells labeled with Tractin-mRuby (magenta) in fluorescently labeled collagen matrix (green) in a multi-well plate over three hours. Scale bars: **A** 20 μm, **B, C** 50μm, inset (from left to right) 20 μm, 10 μm, 20 μm, **D** 100 μm, **E** 20 μm.

To illustrate the large volumetric coverage, we imaged a 4x expanded liver tissue section containing hepatocellular carcinoma, labeled for DNA with SYTOX Green and mitochondria labeled with Hsp60 (**Figure 4B**). After deconvolution, one can discern sub-organelle features of mitochondria, and nucleoli are well resolved also in the axial direction (**Figure 4C**). The height of the imaging stack was ∼130 microns, which is the theoretically estimated remote focusing range (**Table 1** and **Supplementary Figure 1-2**). While we acquired a taller volume with mechanical tiling using the water objective, the air objective allows the acquisition of large volumes via optical scanning (no mechanical motion of the sample or the primary objective).

Traditionally, if an air objective is used to image into a watery sample, aberrations and axial distortions of the imaging volume result from the large refractive index mismatch^14^. We performed comparative imaging with a spinning disk using the same air objective and acquired ground truth data with a water immersion lens (**Supplementary Figure 5**). Importantly, miOPM using an air objective as the primary lens is not subject to the distortions the spinning disk confocal experienced with the same air objective.

Due to the large z-coverage that can be optically accessed, the air objective can be leveraged to rapidly image sufficiently transparent model organisms while avoiding mechanical agitation. In **Figure 4D** and **Supplementary Video 8,** we imaged an amoeba C*haos carolinensis* that was microinjected with Lifeact-GFP2 protein. Within the imaging volume, one can see four vacuoles in the uropod, which is hypothesized to coordinate migration in Amoeba^26,27^.

To highlight the potential for rapid multi-well imaging, we volumetrically imaged cells in 384 wells in four colors (**Figure 1H**). By dispensing with immersion fluid, each well is imaged with similar imaging performance, and in principle, multi-well plates can be rapidly swapped. Besides adherent cells, we also imaged cancer cells in 3D microenvironments. To this end, we seeded SU8686 cell, labeled with Tractin-mRuby, in fluorescent collagen labeled with Alexa Fluor 568 NHS ester in a Cell culture chamber slide with 8 wells. **Figure 4E** and **Supplementary Video 9** shows three selected time points of a cancer cell navigating the collagen network over the course of three hours. Notably, in the path of the cancer cell, collagen fibers appear to be enriched, which has been previously described as a worrying mode of translocation of cancer cells in soft microenvironments.^28^

### Adapting miOPM to cleared tissue imaging

The use of the zoom lens allows miOPM to optimally image samples with refractive indices different from water. After we calibrated the zoom lens by measuring the effective magnification (**Supplementary Figure 3**), we assessed the imaging performance using a channel, decorated with 100 nm diameter fluorescent nanospheres, containing a solution of known refractive index (n=1.47, **Figure 5A**). When the magnification into the remote space was left at 1.33 (as previously used for watery samples), imaging performance was degraded by spherical aberrations at the top of the channel (**Figure 5B**). When we adjusted the zoom lens to a magnification of 1.47, diffraction limited performance was restored (**Figure 5C**). Widefield illumination was used in **Figure 5B-C** to better show the axial features of the detection point spread function.

**Figure 5.**
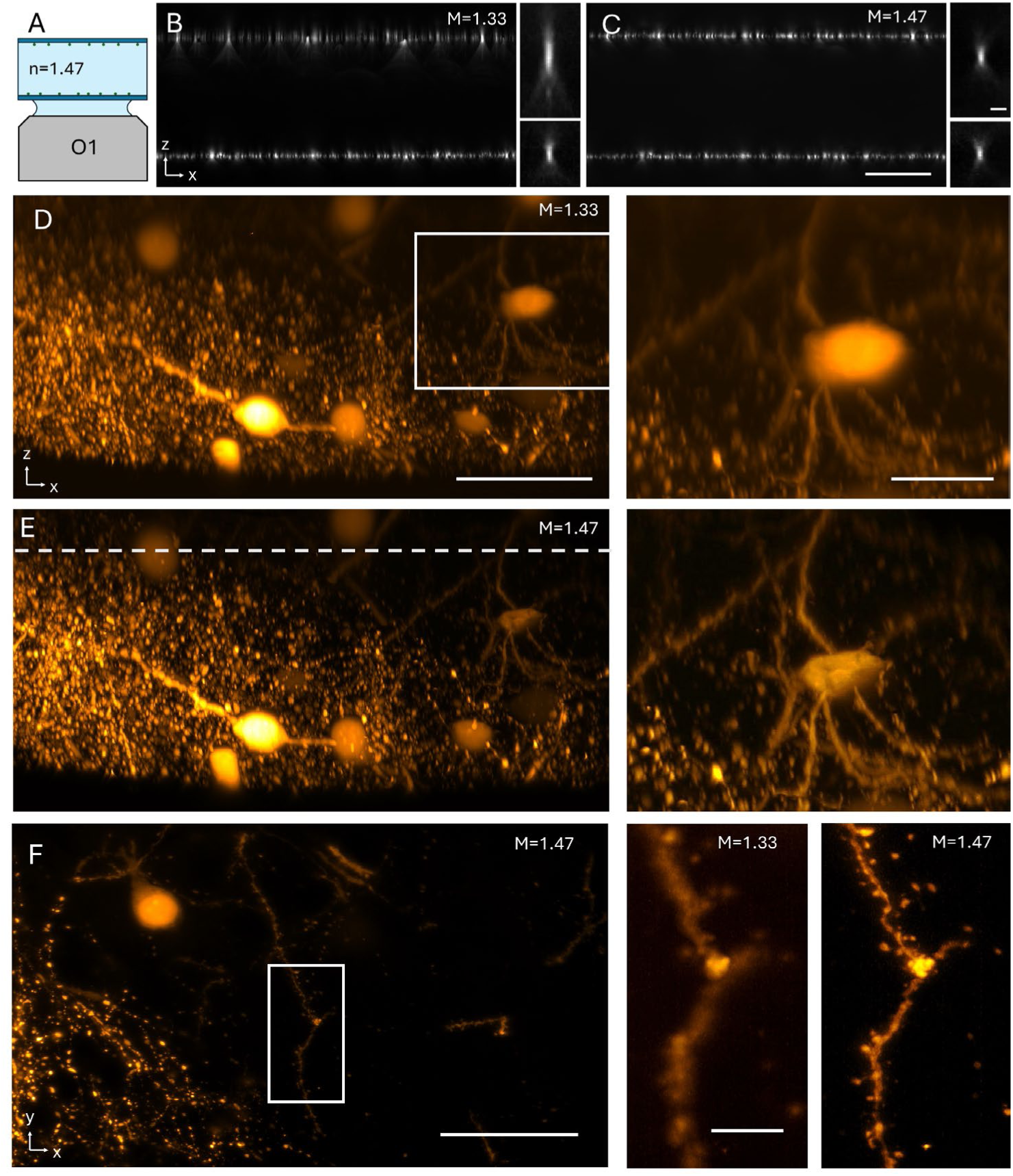
Cleared tissue imaging with refractive index adaptation. **A** Imaging of fluorescent nanospheres in a microchannel containing media of known refractive index of 1.47. **B** Imaging of bead layer at the top and bottom of the channel, with the zoom lens set for water magnification. Widefield illumination was used to show the full extent of the detection PSF. **C** The same volume imaged with the zoom lens set to a magnification of 1.47. **D** Imaging of a Clarity cleared mouse brain section, labeled for Thy1-eGFP with the zoom lens set to 1.33 magnification. **E** The same volume imaged with magnification set to 1.47. **F** X-Y slice along the dotted line in E imaged with 1.33 magnification. **G** Same XY slice but imaged with 1.47 magnification. Scale bars: **C** 20 μm, insert 1 μm; **D** 50 μm, insert 20 μm; **F** 50 μm, insert 10μm.

To show the potential of miOPM for cleared tissues, we imaged a slice of a CLARITY cleared mouse brain labeled with Thy1-eGFP. **Figure 5C** shows an axial cross-section through a volume that was acquired with the zoom lens magnification left at 1.33. This is representative for an OPM system that was optimized watery samples but is used for imaging such a cleared tissue. As it is visible in the magnified inset, imaging quality degrades deeper within the sample.

**Figure 5D** shows the same volume but imaged with the zoom lens set to a magnification of 1.47 to match the refractive index of the sample (measured with a refractometer). As can be seen in the inset, imaging quality remains high throughout the volume, and small features that may represent spines become resolvable even in the axial cross-section.

**Figure 5F-G** shows an X-Y slice, located ∼70 microns above the tissue surface, as imaged with the zoom lens set to 1.33 and 1.47, respectively. The refractive index mismatch in **Figure 5F** obscures fine details, such as spines that can be clearly seen in **Figure 5G**.

## Discussion

We have introduced miOPM, a light-sheet platform that can be adapted to imaging needs in terms of resolution, sensitivity, volumetric coverage, and sample refractive index. We presented applications that range from sensitive imaging of adherent cells with oil immersion objectives, to large volumetric coverage of expanded tissues and model organisms with a water immersion objective and contact-less multi-well imaging with an air objective. This stands in contrast to previously published high-resolution OPM systems, which excelled at imaging adherent cells, but were less suited to image large samples^29,30^. This limitation stems from several factors, including the short remote focusing range for a high-NA OPM systems, and the resulting tight detection PSF, which necessitates the use of thin and short light-sheets. In contrast, OPM systems with intermediate NA (0.8-1.1) enable larger volumetric coverage at the cost of reduced resolution and sensitivity^11,31,32^. With miOPM, users can now balance resolution, volumetric coverage and sensitivity simply by switching the primary objective. While this is a trivial feature on traditional optical microscopes, such as a confocal, this is not the norm in LSFM and OPM in particular.

In this manuscript, we have now rigorously derived the AIRR theory, which enables miOPM to seamlessly use different objectives. We leveraged this capability to perform both sensitive imaging of subcellular dynamics but also imaging multicellular samples with larger volumetric coverage than provided by prior high NA OPM systems. Furthermore, the use of a primary air objective represents an interesting edge case for AIRR, as it is the largest refractive index mismatch between sample and objective presented herein. Importantly, we demonstrated that the theory corresponds well to practice (in terms of volumetric coverage, shift invariance of the PSF and distortion free volumetric imaging). Practically, the air objective also enables multi-well plate imaging, as no immersion media must be maintained between the objective and the plate, enabling high scan speeds, rapid plate swapping and long observations times.

We further demonstrated the potential of miOPM for correlative light and electron microscopy. These correlative datasets establish a bridge between live-cell observations of seeded α-synuclein uptake, and the ultrastructural context of aggregates captured in situ. In previous work, we demonstrated by correlative light and electron tomography that α-synuclein inclusions in human brain tissue recruit membranous organelles^33^. Applying miOPM to a live-cell seeding model may clarify the link the mechanistic process of fibril uptake and templated aggregation in neurons to the pathological inclusions observed in human Lewy bodies, a defining hallmark of Parkinson’s disease and related synucleinopathies^34^. We also envision that miOPM with a primary air objective will be valuable for optical imaging at cryo-temperatures^35^, as it enables distortion free, rapid 3D imaging in this context.

A zoom lens allows miOPM to adapt continuously to a range of refractive indices spanning 1.33 to 1.51, covering a host of tissue clearing and expansion techniques. As demonstrated here, matching the refractive index is paramount for high-resolution 3D imaging, as severe losses in resolving power and non-trivial axial distortions occur otherwise. Since the zoom lens is automated, refractive index tuning can be guided by refractometer measurements or potentially optimized with a smart microscope control software paradigm^36,37^. An adaptable remote focusing system with a zoom lens could also advantageously be implemented in other microscopes, such as a spinning disk^38^ to enable high resolution multi-immersion imaging. Notably, a correction collar cannot perform this type of volumetric correction for refractive index mismatch. Collars correct a fixed amount of spherical aberration experienced at a specific depth for a given medium. Thus, to achieve a similar level of correction would require constant adjustment of the correction collar while acquiring a z-stack, which is impractical for live imaging. Moreover, for an OPM, where the imaging plane is tilted, a single collar correction cannot apply to the entire field of view.

Taken together, miOPM extends the application range of high-resolution OPM platforms and traditional LSFM systems. By unifying versatility with simplicity, we anticipate that miOPM will redefine the standard configuration for OPM platforms. Beyond its immediate applications, miOPM establishes a blueprint for future light-sheet technologies that are adaptable, accessible, and broadly deployable-even in core facility environments with diverse and varied user needs. We envision that the framework presented here will accelerate the democratization of high-resolution LSFM, consolidate fragmented approaches into a single versatile platform, and enable a wider community of researchers to address biological questions previously out of reach.

## Methods

### Imaging Systems

The imaging in this manuscript was performed with two miOPM systems, one at UT Southwestern Medical Center, and one at the Calico Life Sciences LLC.

### UT Southwestern microscope setup

The optical setup of the system at UT Southwestern is shown in **Figure 1C**, which consists of three major units: the remote focusing system, the illumination unit and the tertiary imaging unit.

The main part of remote focusing system comprises the primary objective (O1, Nikon 40X. Options: Air, MRD70470; Water, MRD77410; Silicone Oil, MRD73400; Oil, MRH01401), the secondary objective (O2, Nikon 40X, MRD70470) and two tube lenses (TL1: ITL-200, Thorlabs; TL2: zoom lens, 132.5 to150 mm EFL). Two objectives are placed back-to-back with the tube lenses to ensure the conjugate mapping between their back focal planes (BFPs). It should be noted that the BFP of the objectives are positioned within the lenses, with distances to the lens thread varied between objectives. Therefore, we used customed lens tubes to maintain the BFP of all primary objectives at the same location when exchange. Two galvo mirrors (Saturn 9B, ScannerMAX) are inserted between the tube lenses for fast image space scanning^39^.

The magnification of the remote system is set to match with the refractive index of the samples. Given that we are using the objectives with the same EFL, it is defined by the ratio of tube lenses’ EFL and using a zoom lens as the TL2 allows us to freely adjust the magnification. The zoom lens in this work is a three-element optical system with fixed back and front focal planes^13,40^ .Two examples of lens layout with EFL of 132.5 mm and 150 mm are shown in **Supplementary Figure 2A-B** and **Extended Figure 2**. Lens 1 and Lens 3 are positive lenses, assembled from a pair of 300-mm EFL lenses (47-649, Edmund) and 250-mm lenses (47-647, Edmund) respectively; Lens 2 is a single negative lens with EFL of 150 mm (62-495, Edmund). They are mounted on the motorized stages (DDS100, DDS50, Thorlabs) to adjust their positions. The blue and red lines passing through the back and front focal points indicate that the BFP and FFP remain unchanged across different configurations. We listed the position of the lenses of the configuration we mainly used from this work in **Supplementary Figure 2C**. We calibrated the magnification of the overall imaging system and remote focusing unit under ten different EFL settings of the zoom lens shown as **Supplementary Figure 2D-E**.

The illumination unit was designed based on our previously published module^32^. The key components include a fiber-delivered solid state laser module (OBIS Galaxy with laser modules LS 561nm-80mW, LX 488nm-100mW and LX 640nm-75mW, Coherent Inc) and a collimator (CFC11A-A, Thorlabs) to direct the laser into the system, a Powell lens (LOCP-8.9R20-1.8, LaserLine) to generate the light sheet and a resonant galvo (CRS 4KHz, Cambridge technologies) for shadowing artefacts suppression. the light sheet generated by the illumination unit is coupled through a dichroic mirror (Di03-R405/488/561/635-t1-25×36, Semrock) into the optical train of the OPM. After passing through two tube lenses, it is launched in sample space through the primary objective (O1) with an oblique angle to the coverslip. The angle was chosen to be 40 degrees for oil, silicone oil and water objectives, and 50 degrees for the air objective to obtain the best imaging performance for our applications. However, a smaller tilt angle is possible for the oil and silicone oil primary objectives in principle, to improve collection efficiency and resolution.

Samples are placed on a motorized stage (FTP-2000, ASI) during the imaging. Fluorescence light is detected through the same primary objective. After passing through the remote focusing system, a tertiary imaging system is used to map the fluorescence from the remote space on to a camera to output the images. It consists of a glass-tipped objective (AMS-AGY v2 54-18-9, Applied Scientific instrumentation), a tube lens (EFL= 375, ACT508-750-A x2, Thorlabs) and a scientific CMOS camera (Flash 4, Hamamatsu). We also incorporate an auto-focus unit to keep stable imaging for hour-long acquisition runs in a lab^41^.

The data acquisition computer was a Colfax International ProEdge SXT9800 Workstation. The control software was developed using a 64-bit version of LabView 2016 equipped with the LabView Run-Time Engine, Vision Development Module and Vision Run-Time Module (National Instruments). The software communicated with the camera via the DCAM-API for the Active Silicon Firebird frame-grabber and delivered transistor–transistor logic triggers and analog voltage signals through a field programmable gate array (PCIe 7852R, National Instruments). The control software can be requested from the corresponding authors and will be distributed under a material transfer agreement with the University of Texas Southwestern Medical Center.

### Calico microscope setup

The imaging of the 384 well plate, the organoid and the Amoeba were performed at the Calico Life Sciences LLC. The Calico microscope is a high-throughput (HT) version of the original single-objective light-sheet (SOLS) microscope that was named HT-SOLS, leverages the any immersion remote refocus AIRR technology^13^ and the zoom lens, and optionally uses real time multi-angle projections^42^. The main differences to the system at UT Southwestern are the tilt angles of the tertiary imaging system, the magnification of the tertiary imaging system and the galvo scanning setup.

#### Emission path

The sample is held by an automated XY stage (Physik Instrumente C-867.2U2) and the macroscopic Z location of the XY stage is adjusted with a coarse Z drive (Dual synchronized Thorlabs MLJ motorized vertical translation stages). Enabled by the AIRR concept, a primary objective (O1) is chosen from 3 Nikon 40x options with different numerical apertures (NA) and immersions: NA=0.95 air (MRD00405), NA=1.15 water (MRD77410) or NA=1.3 oil (MRH01401). To maintain the correct Z position for each O1 a separate Z drive (Thorlabs MCM3000 with ZFM2020) is used to automatically adjust the Z of each O1 depending on the choice of immersion (since they have different back focal plane positions). The course Z drive holding the XY stage then used to compensate the O1 Z drive to keep the sample nominally in focus. The O1 objectives are mounted in a high-speed focus piezo with an 800um range (Physik Instrumente E-709.1C1L with P-725.8CDE2) that is then used for all subsequent fine focus adjustments, and is optionally controlled by a hardware autofocus unit (Prior PureFocus850) that sits immediately under the focus piezo. A 2-inch mirror (Thorlabs BB2-E02) folds the light path into the plane of the optical table and is followed by the first tube lens (TL1) which has a 200mm focal length (Thorlabs TTL200). The O1 and TL1 pair produce an intermediate image plane (IP1) with 40x magnification for all choices of O1. A unity magnification scanner images IP1 to IP2 in a lens-galvo-lens configuration using 100mm tube lenses (Thorlabs TTL100-A) and a 10mm diameter galvo (GVS211/M). IP2 is then imaged by the second tube lens (TL2) which in this case consists of the AIRR zoom lens, with a variable focal length in the range 132.5-150mm. TL2 is followed by a dichroic (Chroma ZT405/488/561/640rpc) to couple in the light-sheet and then the secondary objective (O2) which then forms the remote 3D image. O2 is a Nikon 40x NA=0.95 air objective (MRD70470) with a small diameter 170um AR coated coverslip glued to the front (part and service offered by Applied Scientific Instrumentation, SER-GLUE-AR-N40X). The O2 and TL2 pair produce a magnification in the range of 26.5-30x depending on the zoom lens setting, which together with the O1 and TL2 pair gives a remote 3D image magnification in the range of 1.33-1.51x (the intermediate scanner is 1x). The remote 3D image is then collected by a tertiary objective (O3) which is a ‘Snouty’ lens with a 9mm focal length (AMS-AGY v2) and is tilted away from the optical axis of O2 by 45 and 55 degrees for water and air primary objectives, respectively. A filter wheel (Sutter Lambda 10-3) follows O3 to allow different emission filter options (ET445/58M, ET525/50M, ET600/50M, ET706/95M or ZET405/488/561/640m). The tertiary tube lens (TL3) has an effective focal length of 250mm from a custom assembly using a pair of 500mm achromats (Thorlabs ACT508-500-A). The O3 and TL3 pair produce a magnification of ∼27.8x and relay the image to the camera (PCO.edge 4.2 CamLink sCMOS camera). A pair of 20mm galvos (Thorlabs QS20X-AG) sit between TL3 and the camera to perform the optical shearing that is needed for the optional real time multi-angle projection imaging mode.

#### Excitation path

A broadband LED (Thorlabs MBB1L3) sits above O1 and can be used for searching samples with transmitted light. The light-sheet excitation has 4 wavelengths (405nm, 488nm, 561nm and 640nm) from a laser box (Coherent OBIS LS/LX, 1343229) coupled to a beam combiner (Coherent OBIS Galaxy, 1363484) which outputs all 4 laser lines from an FC/APC single-mode polarization-maintaining fiber (NA 0.055). The fiber output is collimated to ∼1.2mm (Thorlabs PAF2-A7A) and levelled to the optical table with a 1-inch mirror (Thorlabs PF10-03-P01). The collimated ∼1.2mm beam is then adjusted with a variable beam expander (Thorlabs BE052-A) to ∼0.8mm and steered by a pair of 1-inch mirrors (Thorlabs BB1-E02) onto a Powell Lens with 30-degree fan angle (Thorlabs LGL130). The fanned light is then collimated by a cylindrical lens with a 24.88mm focal length (Thorlabs LJ1075L1-A). An adjustable slit (Thorlabs VA100CP/M) is placed just before the achromatic cylindrical lens (Thorlabs ACY254-050-A) that generates the light-sheet to modulate the light-sheet thickness. A pair of 1-inch mirrors (Thorlabs BB1-E02 and PF10-03-P01) steer the light-sheet towards the first of three 4f excitation relays (a VA100CP/M could be added between the mirror pair to modulate the light-sheet width in this location). The first relay is for small, automated adjustments to light-sheet tilt and position, and consists of a pair of 100mm achromats (Thorlabs AC254-100-A-ML) bracketing a pair of galvos (Thorlabs GVS202) for tilt, followed immediately by another pair of galvos (Thorlabs GVS202) for position. The second relay is for large manual adjustments to light-sheet tilt and position and consists of a pair of 100mm achromats (Thorlabs AC254-100-A-ML) bracketing a gimbal mirror (Thorlabs GM100/M and BB1-E02) for tilt, followed immediately by another gimbal mirror (Thorlabs GM100/M and BB1-E02) for position. The third and final relay is to couple the light-sheet onto the emission path and consists of a 100mm achromat (Thorlabs AC254-100-A-ML) and the AIRR zoom lens which bracket the dichroic (Chroma ZT405/488/561/640rpc). The AIRR zoom lens and dichroic are shared with the emission path.

### Expanded Liver Specimen Preparation

Hepatocellular carcinoma (HCC) mouse models were generated by Dr. Hao Zhu’s laboratory (UT Southwestern Medical Center). Mice were sacrificed, and livers were harvested and fixed in 4% paraformaldehyde (Thermo Fisher Scientific, #AJ19943K2) overnight at 4 °C with gentle shaking. Fixed liver tissue was then embedded in 2% agarose and sectioned into 100 µm slices for downstream processing. Heat-induced epitope retrieval (HIER) was performed using Tris-EDTA antigen retrieval buffer (Enzo Life Sciences, #ENZ-ACC113) at 95 °C for 30 min. After that, samples were allowed to cool to room temperature for an additional 30 min. Sections were then washed three times in PBS (30 min each) and blocked for 2 h at room temperature in blocking buffer containing 4% bovine serum albumin (BSA; Equitech-Bio, #BAH65) and 0.2% Triton X-100 (Sigma-Aldrich, #X100-100ML) in PBS. Primary antibodies were prepared in blocking buffer: anti-Hsp60 (Abcam, #ab46798, 1:200) and anti-Lamin A/C (Cell Signaling, #4777, 1:100). Tissue sections were incubated with primary antibodies for 24 hat 4 °C with gentle agitation. After incubation, sections were washed three times in PBST (0.1% Tween-20 in PBS; Thermo Fisher Scientific, #BP337500) for 1 h each. Secondary antibodies, goat anti-rabbit IgG (Alexa Fluor™ 568; Thermo Fisher, #A-11036) and goat anti-mouse IgG (Alexa Fluor™ 488; Thermo Fisher, #A-11029), were diluted 1:100 in blocking buffer and applied to samples overnight at 4 °C. After washing in PBST (3 x 1 h), samples were performed with anchoring overnight at room temperature in 0.1 mg/mL Acryloyl-X, SE (Thermo Fisher Scientific) with PBS. Sections were then washed twice for 15 min in PBS and incubated in monomer solution [19% (w/w) sodium acrylate, 10% (w/w) acrylamide, 0.1% (w/w) N,N′-methylenebisacrylamide] overnight at 4 °C. For gelation, samples were gently transferred into a gelation chamber composed of a microscope slide and a silicone spacer (McMaster-Carr, #6459N112) and overlaid with gelation solution [monomer solution plus 0.5% (w/w) ammonium persulfate (APS) and 0.5% (w/w) tetramethylethylenediamine (TEMED)]. Gelation proceeded on ice for 30 min, then for 1.5 h at 37 °C in a humidified incubator. Polymerized samples were denatured in digestion buffer [50 mM Tris (pH 8.0), 1 mM EDTA, 0.5% Triton X-100, 1 M NaCl] containing 8 U/mL Proteinase K (MilliporeSigma, #39450-01-6) at 37 °C for 6 h. After 3 x 30min PBS washes, nuclear staining was performed with SYTOX™ Green (Thermo Fisher, #S7020, 1:1000 in PBS) for 3 h at room temperature, followed by three additional PBS washes (1 h each). Before imaging, samples were expanded in deionized (DI) water for at least 1 h, with three changes of fresh DI water. The estimated expansion factor was ∼4x.

### Multi-well sample preparation

U-2 OS Dmr-PERK G3BP-GFP reporter cells were a gift of Lauren LeBon and Carmela Sidrauski. Cells were grown at 37C/5% CO2 in DMEM supplemented with 10% FBS, L-glutamine and antibiotic-antimycotic. Unless otherwise noted, all liquid handling steps were performed using an Agilent BioTek EL406 washer/dispenser. For imaging studies, 3000 cells were seeded to each well of a 384 well flat-bottomed PhenoPlate (Revvity cat no 6057308) in a volume of 50 ul of growth medium. Cells were allowed to adhere to the plates for 24h, before dosing with small molecule compounds dissolved in DMSO using an Echo 655 acoustic liquid handler. Cells were incubated for 16h with 1, 2, or 5µM of compound. Mitochondria were stained for 30 min at 37C by the addition of 10µl/wl of MitoTracker DeepRed FM (TheroFisher cat no M22426) to a final concentration of 100nM. After mitochondrial staining, cells were fixed for 20 minutes at room temperature by the addition of paraformaldehyde (Electron Microscopy Sciences cat no 15710-S) to a final concentration of 4% w/v. The media-mitotracker-fixative mix was aspirated, and cells were stained for 30 minutes at room temperature in 20µl of 1x Ca/Mg-free HBSS (ThermoFisher cat no 14065-056) supplemented with 0.1% v/v Triton X-100 (ThermoFisher cat no 28314), 1% BSA, 1µg/ml Hoechst 33342 (ThermoFisher cat no H3570), 8µM Phalloidin CF-430 (Biotium cat no 00054), 10µM Revvity PhenoVue 493 (Revvity cat no 00054), and 1.5µg/ml WGA-AlexaFluor 555 (ThermoFisher cat no W32464). Cells were washed 2x with 60µl/wl of Ca/Mg-free HBSS, and stored in 60µl/wl Ca/Mg-free HBSS supplemented with 0.001% w/v sodium azide.

SU8686 Cells were embedded in 2% fluorescence collagen and seeded on a glass-bottom chamber slide (Ibidi, Cat.No:80800). SU8686 pancreatic ductal adenocarcinoma cells were acquired from ATCC, CRL-1837. The cells were labeled with Tractin-mRuby and prepared by our colleague, Gabriel Muhire Gihana. Cells were grown at 37°C/5% CO2 in RPMI supplemented with 10% FBS, L-glutamine, and antibiotic-antimycotic. The preparation of fluorescence collagen was described elsewhere^43^. Briefly, an aliquot of collagen (3mg/mL, PureCol, Type I Bovine Collagen Solution, Advanced BioMatrix, #5005) was covalently labeled with an Alexa Fluor 488 NHS ester. The fluorescence-labeled collagen was mixed with unlabeled collagen at a ratio of 1:9 (total 64%), and then diluted with dH2O (25%), 10X PBS(10%), and 1M NaOH(1%) to reach a final concentration of 2% fluorescence collagen. Cells were mixed with fluorescence collagen, and before collagen polymerization, seeded on the Cell culture chamber slide with 8 wells (Ibidi, Cat.No:80800).

### Preparation of CD8+ T Cells

CD8+ T cells were obtained from blood leukocyte cones purchased from NHS Blood and Transplant, John Radcliffe Hospital, Oxford, UK. Blood cones were used under the ethical guidelines of NHS Blood and Transplant. The Non-Clinical Issue division of National Health Service approved the use of blood leukocyte cones at the University of Oxford (REC 11/H0711/11). Briefly, CD8+ T cells were isolated from blood using RosetteSep™ Human CD8+ T Cell Enrichment Cocktail (Stemcell technologies, cat #15063) followed by EasySep™ Human Naïve CD8+ T Cell Isolation Kit (Stemcell Technologies cat#19258) according to manufacturers protocols. Naive T cells were checked for purity (>97%) using flow cytometry e.g. (CD3+CD45R0-CCR7+). T cells were then activated using anti-CD3/CD28 Dynabeads for 2 days and rested for at least 2 days after removing the beads prior to use.

### Preparation of hTERT RPE-1 Cells

hTERT RPE-1 cells (ATCC CRL-4000) were lentivirally transduced with a modified pLVX-shRNA1 vector in which mNeonGreen–EB3 was placed downstream of a truncated CMV promoter (Addgene #110718). Cells were maintained in ATCC-formulated DMEM/F12 supplemented with 10% FBS and 1% antibiotic–antimycotic (anti-anti) and were routinely screened for mycoplasma. For imaging, cells were plated on Ibidi 35-mm glass-bottom dishes with a #1.5 coverslip and exchanged into Gibco FluoroBrite DMEM to reduce background fluorescence.

### Preparation of CLARITY cleared Thy1-eGFP mouse brain sample

Thy1-eGFP mouse (Tg(Thy1-EGFP) MJrs/J, JAX Strain 007788), 8 weeks old, was used for CLARITY tissue clearing. The mouse was transcardially perfused with 4% paraformaldehyde (PFA, 15710-S, Electron Microscopy Sciences), and the brain was extracted and fixed in 4% PFA overnight at 4°C.

The PFA-fixed brain underwent passive CLARITY clearing^44^. The brain was incubated overnight at 4°C in hydrogel monomer (HM) solution containing 1% acrylamide (1610140, Bio-Rad), 0.05% bisacrylamide (1610142, Bio-Rad), 4% PFA, 1× PBS, deionized water, and 0.25% thermal initiator (VA-044, Fisher Scientific). Following incubation, the solution was degassed by removing oxygen and replacing it with nitrogen using a vacuum pump. The sealed sample was polymerized at 37°C for 6 hours. The polymerized brain was sectioned into 1mm thick coronal sections using a vibratome.

Brain sections were passively cleared in sodium dodecyl sulfate (SDS)/borate clearing (SBC) buffer (4% SDS, 0837, Amresco; 0.2 M boric acid, B7901, Sigma-Aldrich, in deionized water, pH 8.5) at 37°C with gentle shaking. Clearing progress was monitored daily until samples became transparent, typically around 1 week for 1mm sections. Cleared samples were washed in 0.2 M boric acid (pH 8.5). Prior to imaging, samples were refractive index-matched using RapiClear (SunJin Lab, RI 1.47).

### Preparation of synuclein-treated neurons

Primary neuronal culture. Striatal neurons were isolated from embryonic day 18 C57BL/6 mouse embryos and plated directly onto Quantifoil holey carbon EM grids. Neurons were maintained in Neurobasal medium supplemented with B27 and GlutaMAX and cultured for 14 days in vitro (DIV14), at which stage they form mature axonal and dendritic networks suitable for correlating dynamic molecular events with ultrastructure.

Preparation and labeling of α-synuclein fibrils. Recombinant human α-synuclein was purified, assembled into pre-formed fibrils (PFFs), and sonicated to generate seeding-competent fibrils. For fluorescence tracking, PFFs were covalently labeled with Alexa Fluor 647 (aSyn-647).

Fluorescence imaging with miOPM. Prior to fibril addition, neuronal plasma membranes were labeled with MemGlow-488 (Cytoskeleton, Inc.) to delineate overall architecture, including thin axonal and dendritic processes. Labeled aSyn-647 PFFs were then introduced to the culture medium at a final concentration of 100 nM. Live fluorescence imaging was performed on a custom-built multi-immersion oblique plane microscopy (miOPM) system. Imaging revealed discrete aSyn-647 puncta along axons, consistent with internalized clusters.

Cryo-EM sample preparation and imaging. Immediately after light microscopy, grids were manually blotted from the reverse side and plunge-frozen in liquid ethane cooled by liquid nitrogen to vitrify neuronal ultrastructure. Frozen grids were first screened on a Talos Arctica (Thermo Fisher Scientific, 200 kV) to assess ice thickness and sample distribution. High-resolution cryo-EM was subsequently performed on a Titan Krios (Thermo Fisher Scientific, 300 kV) equipped with a K3 direct electron detector (Gatan).

Correlation. Grid square registration was used to align fluorescence and cryo-EM data. Overlays confirmed that fluorescent aSyn-647 puncta corresponded to aggregates within neuronal processes, establishing the ability of miOPM-enabled CLEM to connect dynamic fluorescence readouts with cryo-preserved ultrastructure.

### Alveolar organoid preparation

Alveolar epithelial type II (AT2) cells were isolated by dissociating the lungs of female C57BL6/J mice (The Jackson Laboratory, strain #000664) for the culture of alveolar organoids. Briefly, lungs were inflated intratracheally with Dispase (Stem Cell Technologies #07913) and DNase I (Sigma #D5025) and dissociated in gentleMACS C tubes (Miltenyi Biotec #130-093-237) on a gentleMACS Dissociator (Miltenyi Biotec #130-134-029) in DMEM (Corning #10-013-CV). AT2 cells were enriched via magnetic depletion of immune and endothelial cells (Stem Cell Technologies #19860), then stained using anti-EpCAM and MHCII antibodies (Biolegend #118230; #107670) and FACS sorted by gating on the EpCAM^+^ MHCII^+^ population. Isolated mouse AT2 cells were then seeded in growth factor reduced Matrigel (Corning #354230) and grown in a cell culture incubator at 37°C with 5% CO_2_ in AMM media (Katsura et al., 2020 PMID 33128895).

For immunostaining, alveolar organoids were centrifuged at 1900 *g* for 10 min then fixed in 4% PFA (Electron Microscopy Sciences, #15714) at room temperature for 45 min. Fixed organoids were permeabilized in 0.5% Triton X-100 at room temperature for 1 hour, then blocked in 3% normal goat serum with 0.1% Triton X-100 at room temperature for 1 hour. Cells were stained in Muc1, RAGE, and TOM20 antibodies at 4°C overnight (Thermo #MA511202; R&D Systems #MAB1179; Proteintech #11802-1-AP, respectively). Organoids were washed three times with PBS, then stained with secondary antibodies at 4°C overnight (Jackson Immunoresearch #127-545-099, Thermo #A11077, Thermo #A32733). Samples were stained with DAPI (Thermo #D1306) at room temperature for 20 minutes, then washed three times in PBS before imaging in PBS by OPM with a 40x water objective.

### Labeling and visualization of mitochondria and mtDNA

U-2 OS or HeLa cells were cultured in DMEM (D5796; Sigma-Aldrich) that was supplemented with 10% FBS, 25 mM HEPES, and 1% penicillin/streptomycin. To label mitochondrial DNA, HeLa cells were passaged to glass-bottom dishes (D35-14-1.5-N; CellVis), stained with 25nM MitoTracker CMXRos (M7512; Thermo Fisher Scientific) and 1:1,000 PicoGreen (P7589; Thermo Fisher Scientific) in culture media for 1 hour at 37°C, washed twice, and imaged live. To visualize the mitochondrial outer membrane, U-2 OS cells were transiently transfected with GFP-OMP25 (Addgene 141150, a kind gift of Gia Voeltz) with Lipofectamine 3000 (Thermo Fisher Scientific) according to manufacturer’s instructions, cells were passaged to glass-bottom dishes, allowed to adhere, and imaged live.

Data post processing

Microscope raw data was de-skewed and rotated using the PetaKit5D^45^. On some datasets, deconvolution using the two step OMW method was applied. The following datasets are deconvolved: **Figure 1F&G, Figure 2A-E, Figure 3E-F and Figure 4B-C & E**. The rest of the data is shown without deconvolution.

### Particle Detection, Tracking, and Visualization

Deconvolved, sheared, and rotated image volumes were cropped to isolate individual cells of interest. Particle detection and tracking were performed using uTrack-3D. Particles were identified using the bandPass watershed algorithm with an intensity threshold of 30 and low and high-pass frequency cutoffs of 2 and 10, respectively. For trajectory linking, the maximum gap closure was set to 3 frames, and both merging and splitting events were permitted. Detection and tracking results were exported to CSV and imported into Python for visualization with napari. DNA and mitochondria were visualized using green and magenta colormaps, respectively, with intensity ranges set to [0, 980] and [300, 3000]. Both channels were displayed with isotropic scaling and kaiser interpolation. Detected particles were rendered as semi-transparent, 2-pixel points, with face color mapped to amplitude and white borders to enhance visibility. Track trajectories were displayed with a tail length of 20 timepoints and tail width of 2. Representative frames from distinct view angles and zoom levels were exported as PNG files and assembled into movies using Fiji.

## Acknowledgements

We thank Dr. Marcel Mettlen for providing the fixed ARPE cell and Dr. Hao Zhu for the liver tissue samples. We are grateful for in-depth discussion on remote focusing with James Manton and Alex Corbett. We are also thankful to Dr. Daniel Gottschling and Dr. Julia Lazaari Dean for feedback on the manuscript. This work was funded by National Institutes of Health: National Institute of General Medical Sciences (R35GM133522 to R.F., R35GM119768 to M.H., R35GM137894 to J.R.F., RM1GM145399 to K.M.D.); National Institute of Biomedical Imaging and Bioengineering (R01EB035538 to R.F.); National Institute of Mental Health (DP2MH119423 to R.T.); National Cancer Institute (U54CA268072 to K.M.D. and R.F.).

## Data availability

The raw data underlying this manuscript will be shared in a public repository when the manuscript is accepted in its final form.

## Code availability

All algorithms, code and software used in this study are publicly available (peer reviewed or preprint)

## Extended Figures

**Extended Figure 1.**
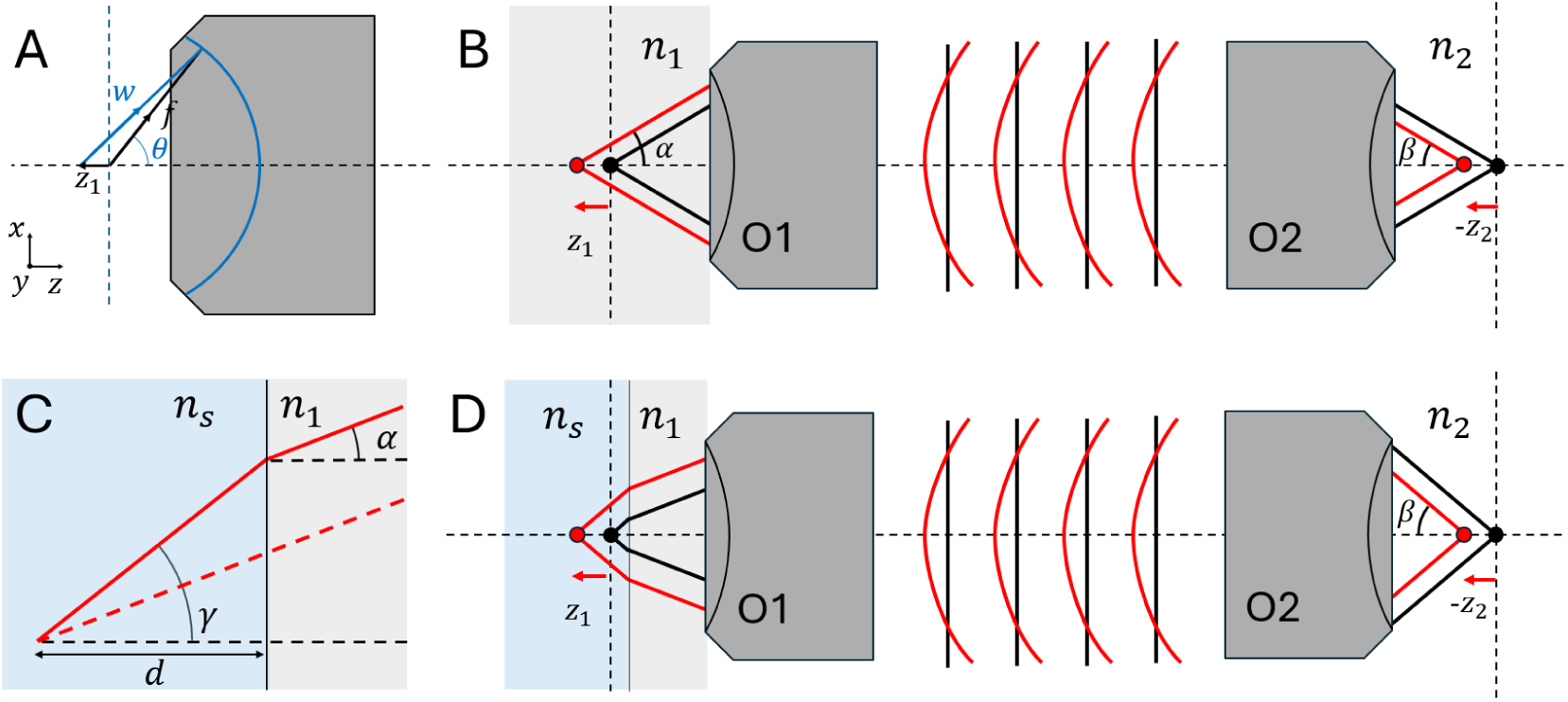
Principle of remote focusing (B) and Any Immersion remote refocusing (AIRR, D). **A** Phase introduced by the axial displacement of a point within a high-NA objective. **B** Conventional remote focusing imaging system. FP (vertical dashed lines) presents the focal plane of objectives. **C** Phase introduced by sample-immersion media refractive index mismatch. The vertical solid line presents the interface. **D** Remote focusing imaging system with refractive index mismatch. When the mismatch interface is on the focal plane, we can get the largest imaging range in depth.

**Extended Figure 2.**
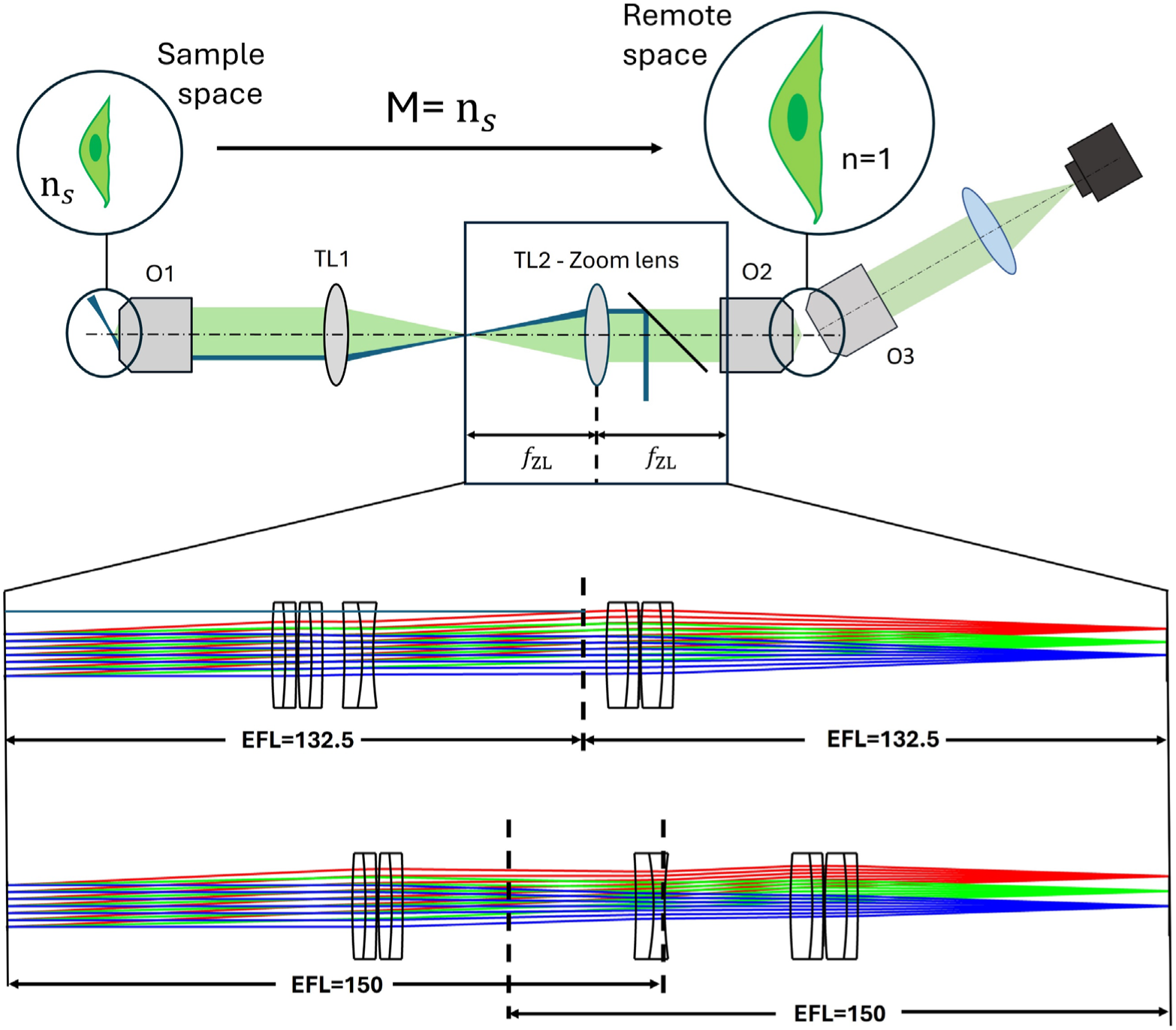
Zoom lens system to adapt miOPM to different sample media. To minimize aberrations and distortions, the magnification into the remote space needs to equal the refractive index of the sample (when using an air objective for O2) .The relay lenses in our OPM system consist of a tube lens (TL1, EFL 200) and the custom zoom lens with a variable focal length from 132.5 to 150mm (ray tracing for two configurations shown below). O1 and O2 have the same EFL (8mm), thus the magnification M to the remote space between O2 and O3 can be continuously varied from 1.51 to 1.33X. The vertical dashed lines indicate principal planes of the Zoom lens system at EFL=132.5 mm and EFL=150 mm, respectively.

