## Supplementary Information for "Multi-immersion Oblique Plane Microscope (miOPM): A reconfigurable platform for high-resolution Light-Sheet Fluorescence Microscopy"

### Supplementary Notes

#### Supplementary Note 1

E.J. Botcherby et al. introduced the theory of remote focusing and derived the equation to calculate the imaging depth as the depth where the Strehl ratio is above 0.8[1]. However, to simplify their analysis, they treated the system as an imaging system with unit magnification, without accounting for the refractive index mismatch of the sample. Here, we followed their general pupil function to theoretically analyze the imaging depth of remote focusing, considering non-unity magnification and the effects of refractive-index mismatch.

Since most objectives are designed to satisfy the sine condition, and our focus here is solely on calculating the axial imaging range (assuming a point on the optical axis with x=0 and y=0 coordinates), we rewrite the relative path length difference (PD) between the imaging point and the focal point in the first imaging unit (shown as **Extended Figure 1A**) as follows

$$PD = |\boldsymbol{w}| - f = f\left\{1 - \frac{2}{f} z_1 \cos\theta + \frac{{z_1}^2}{f^2}\right\}^{1/2} - f, \quad (1)$$

where $\boldsymbol{w}$ is a vector from a point source with a displacement $z_1$ to the focal point, $f$ is the focal length, $\theta$ is the polar angle in the spherical polar coordinate system and $z_1$ is the amount of axial point source displacement. Replacing $\theta$ with normalized pupil radius $\rho$ and the maximum semi-aperture angle of the lens $\alpha$, we can have the phase introduced by PD as

$$\Delta\Psi_1 = -n_1 k f [\left\{1 - \frac{2z_1}{f}(1 - \rho^2 sin^2\,\alpha)^{\frac{1}{2}} + \frac{{z_1}^2}{f^2}\right\}^{1/2} - 1]. \quad (2)$$

where $k$ is the vacuum wavenumber and $n_1$ is the refractive index of the first objective. Expanding this equation to the term of ${z_1}^2$, we have

$$\Delta\Psi_1 = -\frac{k n_1 \rho^2 \sin^2\alpha}{2f} {z_1}^2 + n_1 k z_1 \sin\alpha\,(\csc^2\alpha - \rho^2)^{1/2}. \quad (3)$$

In the remote focusing system, a second imaging unit is placed back-to-back to the first unit (shown as **Extended Figure 1B**). Then we can write the term from the second imaging unit in the same way as

$$\Delta\Psi_2 = -\frac{k n_2 \rho^2 \sin^2\beta}{2f} {z_2}^2 - k n_2 z_2 \sin\beta\,(\csc^2\beta - \rho^2)^{1/2}, \quad (4)$$

where $\beta$ denotes the maximum semi-aperture angle of the lens 2, whose actual value is affected by $\alpha$ in the first imaging unit (it can be smaller than the physical semi-open angle of L2, and when it's bigger than the lens can allow, it will affect $\alpha$). It is important to note that the displacement $z_2$ has a negative sign in the equation due to the back-to-back configuration of the two imaging units. The focal length $f$ remains the same in both equations, as the objective in our system has the same focal lengths. Then we also have $n_1 \sin\alpha = M n_2 \sin\beta$, where M is the lateral magnification of the whole remote focusing system.

The phase introduced by the remote focusing system will be

$$\Delta\Psi_r = \Delta\Psi_1 + \Delta\Psi_2$$
$$= -\frac{kn_1\rho^2 \, sin^2\,\alpha}{2f}\left({z_1}^2 + \frac{n_1{z_2}^2}{n_2M^2}\right) + kn_1 \, sin\,\alpha \;\left[z_1(csc^2\,\alpha - \rho^2)^{1/2} - z_2\frac{1}{M}(csc^2\,\beta - \rho^2)^{1/2}\right]. \quad (5)$$

When imaging into a refractive-index-mismatched sample (shown as **Extended Figure 1C**), there's additional phase generated, we write it down according to Booth et al[2]

$$\Delta\Psi_s = -kn_1 d \sin\alpha\,[(\csc^2\gamma - \rho^2)^{1/2} - (\csc^2\alpha - \rho^2)^{1/2}], \quad (6)$$

where $\gamma$ is the corresponding angle of $\alpha$ in the sample, $d$ is the depth of the focus into the sample. Because of Snell's law, we have $n_1 \sin\alpha = n_s \sin\gamma$.

Then the overall phase introduced by the remote focusing system and sample (shown as **Extended Figure 1D**) is

$$\Delta\Psi = \Delta\Psi_s + \Delta\Psi_r. \quad (7)$$

We can notice when the magnification matches with the refractive index of the sample ($\mathrm{n}_s$) to air ($\mathrm{n}_2$), $M = \mathrm{n}_s/\mathrm{n}_2$, and when the interface between sample and media of first objective is placed at the focal plane ($d = -z_1$), the second term of $\Delta\Psi_s$ will be canceled by $\Delta\Psi_{sr}$, and the $\Delta\Psi$ can be simplifed as

$$\Delta\Psi = -\frac{kn_1\rho^2\sin^2\alpha}{2f}{z_1}^2\left(1 + \frac{n_1}{n_2}\right), \quad (8)$$

Considering the imaging space is in the sample with the refractive index of $\mathrm{n}_s$, we rewrite it as

$$\Delta\Psi = -\frac{kn_s\rho^2\sin^2\gamma}{f}{z_1}^2 \cdot \frac{1}{2}\left(\frac{n_s}{n_1} + \frac{n_s}{n_2}\right). \quad (9)$$

We also noticed from Eq.(5)-(7) that when the magnigication does not match with the ratio of refractive index, there will be extra terms that decreases the imaging depth, although the phase will still be zero at focal plane ($z_1 = 0$). However, if the interface is not located at the focal plane ($d \neq -z_1$), an additional phase and resulting aberration are introduced to the whole volume, including even the focal plane. We observed this phenomenon experimentally.

Before using Strehl ratio to quantify the imaging rage in z dimention, we need to remove the modes that will not affect the focus point, namely piston ($\Psi_p$) amd high-NA defocus ($\Psi_d$). The phase of high-NA defocus term is $n_s k(1 - \rho^2\sin^2\gamma)^{1/2}$, but in order to make sure its integral is zero, we added a constant C such that

$$\int [n_s k(1 - \rho^2\sin^2\gamma)^{1/2} + C]\,\rho d\rho = 0. \quad (10)$$

Then we can have the form of them as

$$\Psi_p = 1, \quad (11)$$

$$\Psi_d = n_s k(1-\rho^2 \sin^2\gamma)^{1/2} - \frac{2n_s k}{3\sin^2\gamma}(1-\cos^3\gamma), \tag{12}$$

where the second term in $\Psi_d$ is C. Because these modes are orthognal, we can get the coefficients of pistion ($a$) and high-NA defocus ($\delta z$) using dot product

$$a = \frac{\int \Delta\Psi\Psi_p \rho d\rho}{\int(\Psi_p)^2 \rho d\rho} = -\frac{n_s k {z_1}^2 \sin^2\gamma}{2f} \cdot \frac{1}{2}\left(\frac{n_s}{n_1} + \frac{n_s}{n_2}\right), \tag{13}$$

$$\delta z = \frac{\int \Delta\Psi\Psi_d \rho d\rho}{\int(\Psi_d)^2 \rho d\rho} = \frac{12{z_1}^2\cos^2\frac{\gamma}{2}(3+6\cos\gamma+\cos 2\gamma)}{5f(3+8\cos\gamma+\cos 2\gamma)} \cdot \frac{1}{2}\left(\frac{n_s}{n_1} + \frac{n_s}{n_2}\right). \tag{14}$$

Then we can substract the these modes to get the phase

$$\Delta\Psi' = \Delta\Psi - a\Psi_p - \delta z\Psi_d. \tag{15}$$

Now we can insert the phase into the equation of Strehl ratio

$$\begin{aligned} S &= \left|\frac{1}{\pi}\int \exp(j\Psi)\,\rho d\rho d\phi\right|^2 \approx 1 - \frac{1}{\pi}\int (\Delta\Psi)^2 \rho d\rho d\phi \\ &= 1 - \frac{4{n_s}^2 k^2 {z_1}^4\,(3+16\,cos\,\gamma + cos\,2\gamma) sin^8\left(\frac{\gamma}{2}\right)}{75f^2(3+8\,cos\,\gamma + cos\,2\gamma)} \cdot \left(\frac{1}{2}\left(\frac{n_s}{n_1} + \frac{n_s}{n_2}\right)\right)^2. \end{aligned} \tag{16}$$

We numerically calcuted the value of $S$ with $z_1$ and defined the theoretical imaging depth when $S$ reaches 0.8. The results with different objectives are listed in the **Table 1**. The codes for numerical calculations for effective NA based solid angle and imaging depth are provided.

### Supplementary Figures

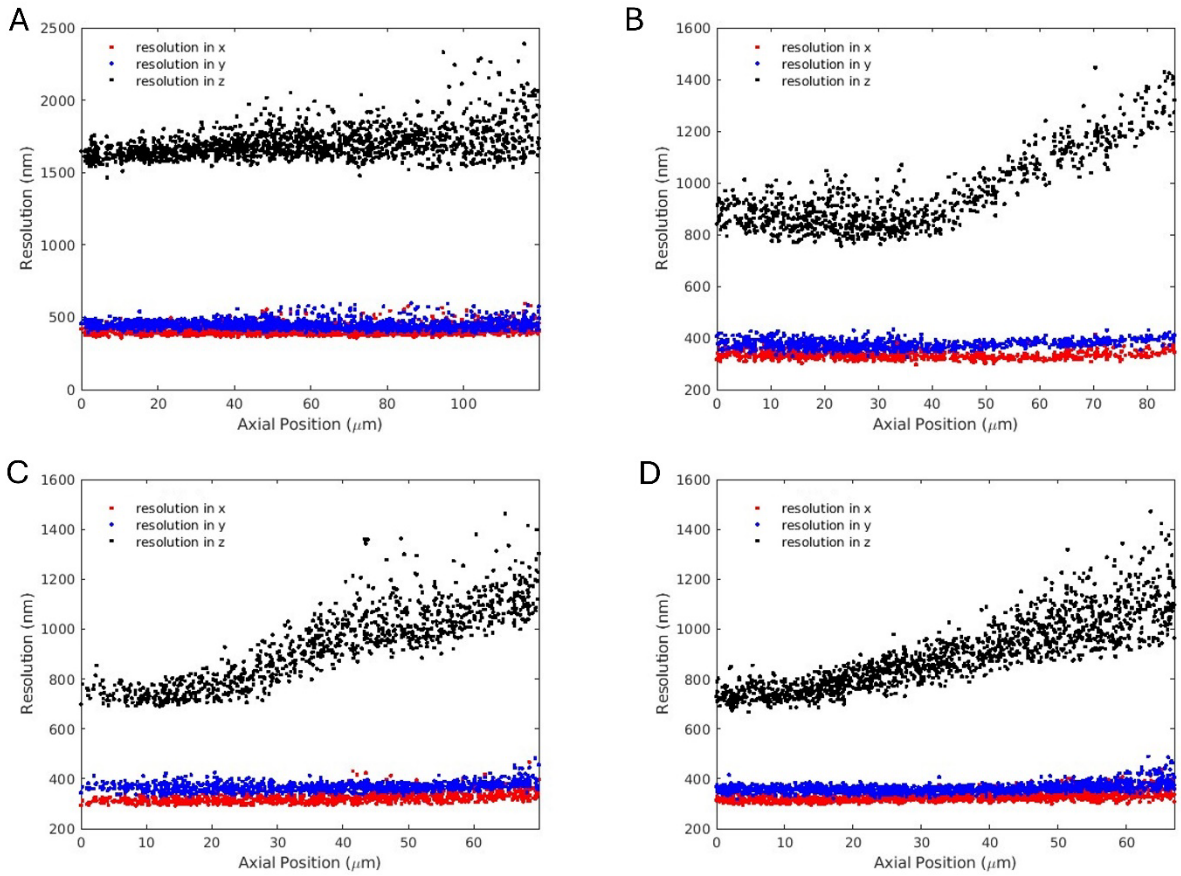


**Supplementary Figure 1. Spatial resolution over depth.** Resolution, as measured by full width half maximum (FWHM) of 100 nm fluorescent nanospheres in Agarose**. A** 40X NA 0.95 Air objective**, B** 40X NA 1.15 water objective, **C** 40X NA 1.25 Silicone oil objective**, D** 40 X NA 1.3 oil objective

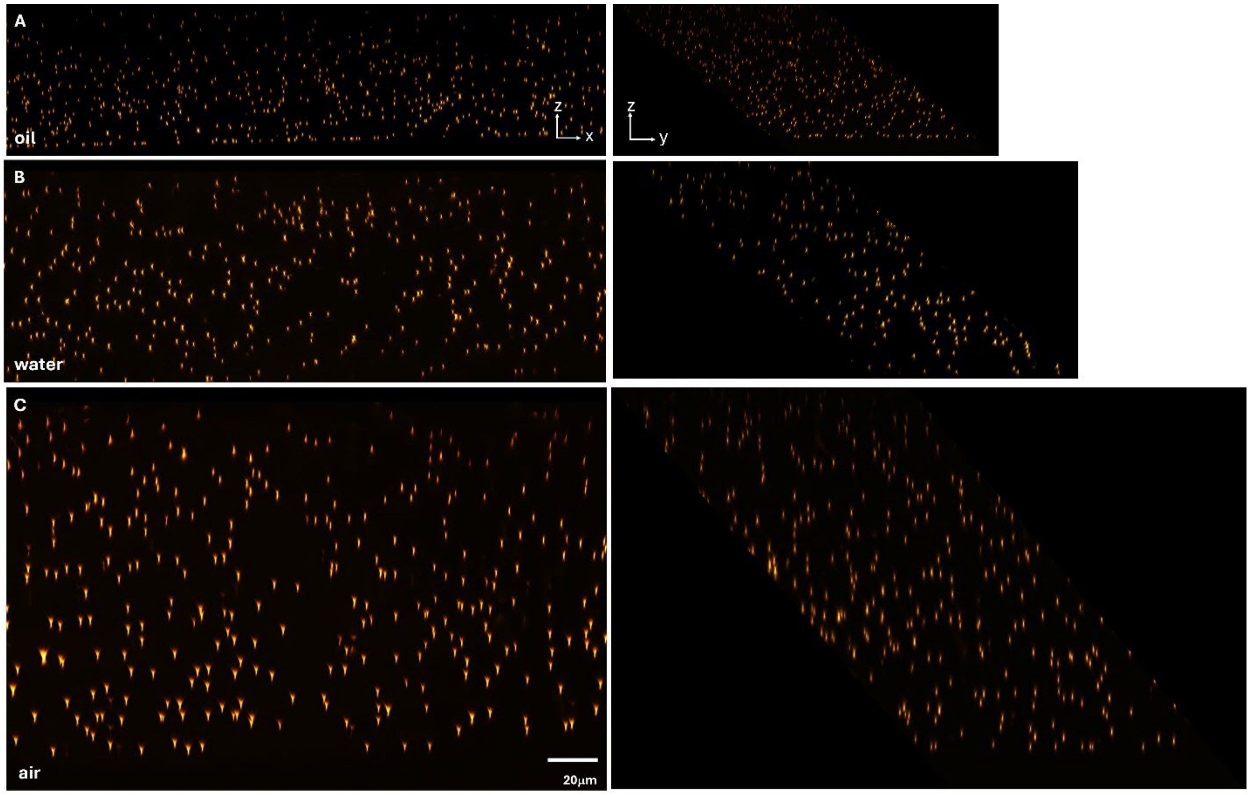


**Supplementary Figure 2. Imaging 100 nm fluorescent nanospheres in Agarose with different primary objectives. A 40** X NA 1.3 oil objective **B** 40X NA 1.15 water objective, **C** 40X NA 0.95 Air objective**,**

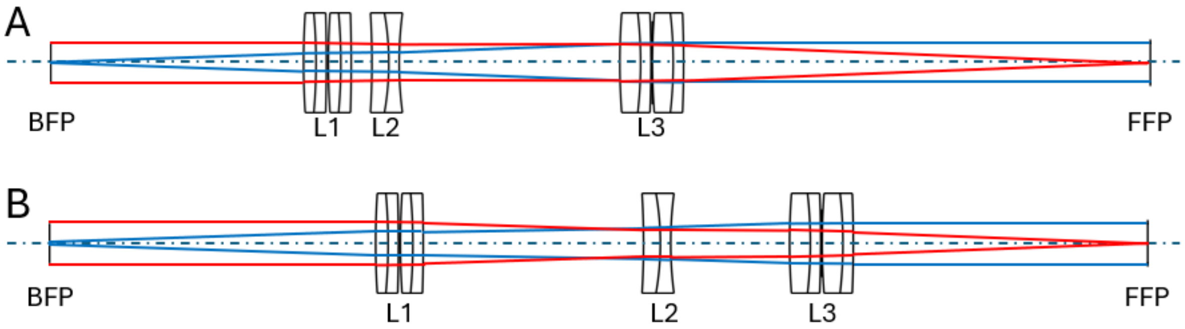


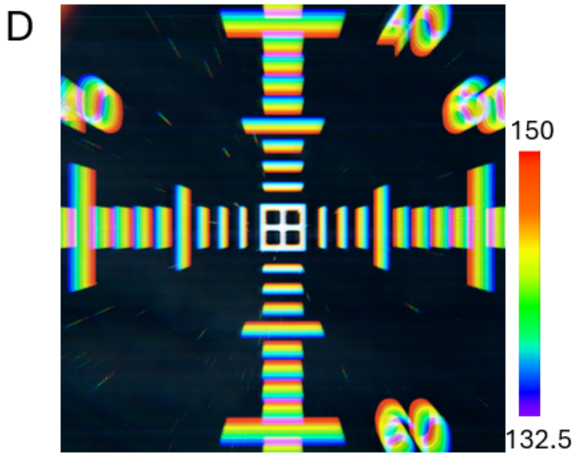


C

| EFL | L1 | L2 | L3 | n |
|---|---|---|---|---|
| 132.5 | 60.90 | 77.36 | 137.06 | 1.51 |
| 135.8 | 65.27 | 89.47 | 143.12 | 1.47 |
| 142.8 | 71.92 | 114.32 | 158.36 | 1.40 |
| 150.0 | 78.62 | 143.25 | 178.49 | 1.33 |

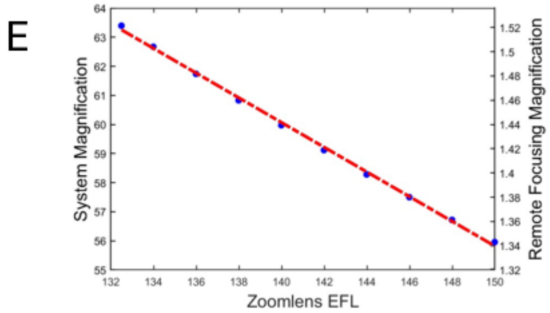


**Supplementary Figure 3. Zoom lens and its magnification calibration.** Optical layout of zoom lens with the EFL of 132.5 mm (**A**) and 150 mm (**B**). BFP, back focal plane; FFP, front focal plane, L1-L3, Lens 1 to Lens 3. The red and blue lines represent the rays focused on the BFP and FFP. **C** Examples of lenses' position with different EFL used in this work. L1-L3 are the distance between each lens and the BFP. The dimensions of EFL and L1-3 are in millimeters. n is the theorical refractive index the system can match. **D** **I**maging stage calibration micrometer on the miOPM microscope under ten different EFL settings of the zoom lens. **E** Measured system magnification (overall magnification of miOPM) and magnification into the remote space as a function of the EFL settings of the zoom lens. A linear fit is shown in red.

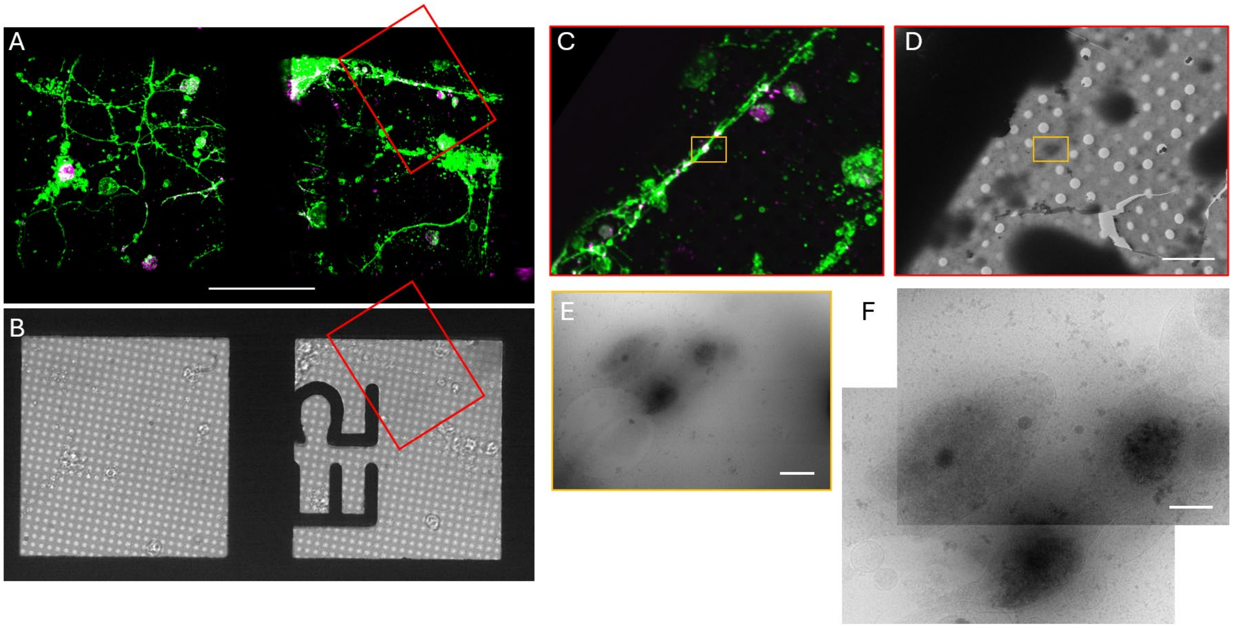


**Supplementary Figure 4. Correlative light and electron microscopy. A** Fluorescence imaging of striatal neurons (green) and alpha-synuclein (magenta) cultured on an EM copper grid as imaged by miOPM. **B** Wide-field imaging of the same region in A, acquired by the wide-filed imaging unit installed in miOPM. **C** The boxed region in A. **D** Wide-field image of the same region in C after plunge-freezing, as imaged by wide-field imaging unit in cryo-EM. **E** The boxed region in **D**, as imaged with higher magnification. **F** Cryo-EM imaging of the same region as E. Scale bars: **A** 50 um, **D** 10 um, **E** 1 um, **F** 500 nm.

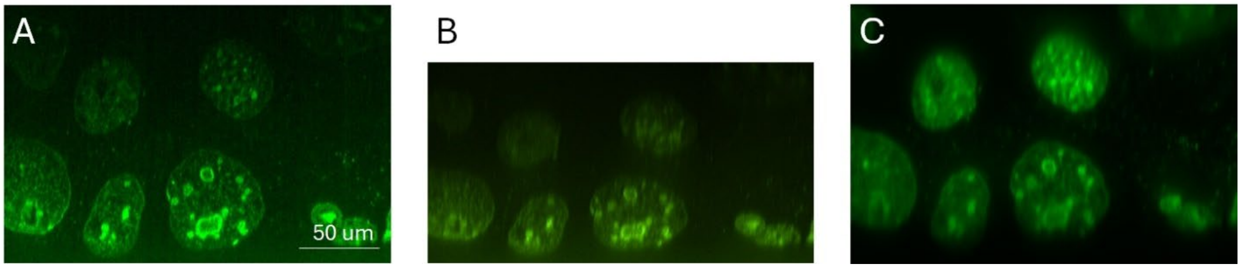


**Supplementary Figure 5. Comparison between confocal and miOPM imaging. A** Cross-section of an image stack acquired with a Spinning disk confocal microscope using a 40X NA 1.15 water immersion objective. The sample is an expanded liver tissue slice labeled with sytoxGreen. **B** Same region, but imaged with spinning disk confocal microscope using an 40X NA 0.95 air objective. **C** Same are imaged with miOPM using an 40X NA 0.95 air primary objective.

### Supplementary Tables

| Figure | Microscope | Primary Objective | tilt angle ° /zoom lens | Acquisition time | Exposure time | Volume size (xyz) um^3 | Pixel size | Laser Power |
|---|---|---|---|---|---|---|---|---|
| 1E | miOPM | Oil<br>Water<br>Air | 40 /1.33<br>40/1.33<br>50/1.33 | N/A<br>N/A<br>N/A | 50ms<br>50ms<br>50ms | 196x241x77<br>226x241x105<br>351x241x162 | 118nm<br>118nm<br>118nm | 42 μW @488<br>32 μW @488<br>25 μW @488 |
| 1F | miOPM | Oil | 40/1.33 | 10s | 10ms | 121x90x38 | 118nm | 28μW@488<br>50μW@561 |
| 1G | miOPM | Silicone Oil | 40/1.47 | N/A | 50ms | 219x202x110 | 107nm | 84 μW @488 |
| 1H | Calico | Air | 55/1.33 | 6.3s | 1ms | 262x210x44 | 175nm | 20, 10, 20, 60( '405', '488', '561', '640') |
| 2A | miOPM | Oil | 40/1.33 | N/A | 50ms | 146x241x38 | 118nm | 0.34 mW@488 |
| 2B | miOPM | Oil | 40/1.33 | 1s | 7ms | 217x236x96 | 122nm | 0.4mW@561 |
| 2C-E | miOPM | Oil | 40/1.33 | 12s | 60ms | 102x86x16 | 118nm | 2.8μW@488<br>5.0μW@561 |
| 2F | miOPM | Oil | 40/1.33 | 7.5s | 36ms | 91x91x21 | 118nm | 0.11mW@488 |
| 2G | Field Synthesis | water | N.A. | 7.5s | 45ms | 83x96x21 | 104nm | 51.5μW @488 |
| 3A | miOPM | water | 40/1.33 | N/A | 20ms | 240x217x597 | 118nm | 22μW@488 |
| 3C | Calico | water | 55/1.33 | N/A | 1ms | 263x267x26 | 175nm | 16, 20, 20, 20( '405', '488', '561', '640') |
| 3E-F | miOPM | water | 40/1.33 | 19s | 50ms | 250x127x17 | 122nm | 80μW@488<br>0.85mW@641 |
| 3G | cryo | NA | 40/1.33 | NA | NA | NA | NA | NA |
| 4A | miOPM | Air | 50/1.33 | 2.74s | 50ms | 85x120x87 | 118nm | 0.3 mW @ 488 |
| 4B-C | miOPM | Air | 50/1.33 | N/A | 50ms | 366x241x162 | 118nm | 0.19mW@488<br>0.37mW@561<br>0.62mW@641 |
| 4D | Calico | Air | 55/1.38 | 4s | 0.5ms | 255x247x109 | 170 nm | 2 @488 |
| 4E | miOPM | Air | 50/1.33 | 16.5s | 25ms | 118x127x68 | 147nm | 24μW@488<br>0.15mW@561 |
| 5B | miOPM | Silicone Oil | 40/1.33 | N/A | 50ms | 56x100x110 | 118nm | 84μW@488 |
| 5C | miOPM | Silicone Oil | 40/1.47 | N/A | 50ms | 56x100x110 | 107nm | 84μW@488 |

| | | | | | | | | |
|---|---|---|---|---|---|---|---|---|
| 5D | miOPM | Silicone Oil | 40/1.33 | N/A | 50ms | 219x117x125 | 118nm | 84μW@488 |
| 5E | miOPM | Silicone Oil | 40/1.47 | N/A | 50ms | 219x117x125 | 107nm | 84μW@488 |

**Supplementary Table 1 Image acquisition parameters.** Laser power in red is the direct output from the laser box in  mW, else laser power is measured in the pupil of the primary objective. Tilt angle is the angle of the tertiary objective and zoom lens refers to the magnification the zoom lens was adjusted to.

### Supplementary Videos

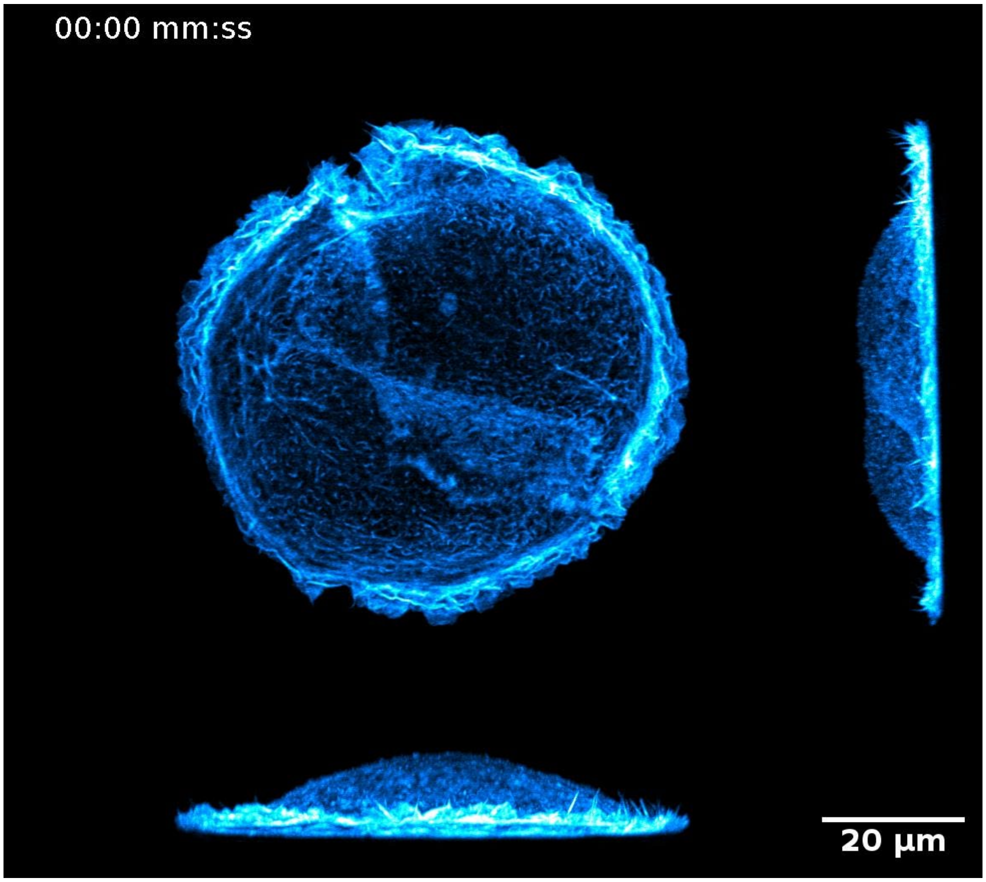


**Supplementary Video 1.** SU8686 cell labeled with Tractin-mRuby as imaged by OPM with an oil immersion objective over 60 timepoints with a 10-s interval. The left and bottom show cross-sectional side views of the cell.

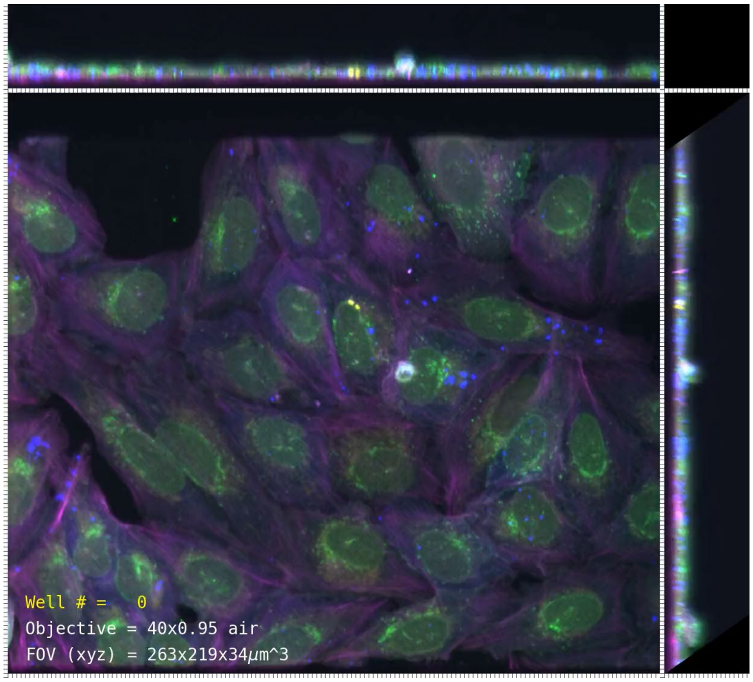


**Supplementary Video 2.** U-2 OS cells stained with Mitotracker Far Red, Hoechst, Phalloidin CF430, Phenovue 493, and WGA-Alexa555, contained in a contact-less multi-well plate imaged with an air objective on miOPM. 384 wells were acquired with a 6.3-s time interval.

**Supplementary Video 3**

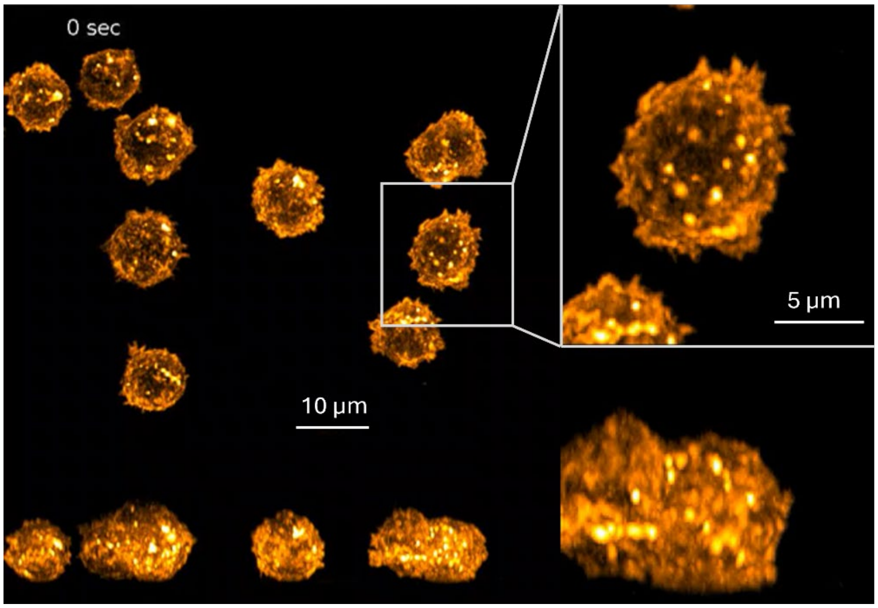


**Supplementary Video 3.** CD8+ T-cells, membrane labelled with CellMask Orange, as imaged with the miOPM. On the right, magnified versions of the boxed regions on the left are shown. White arrows point at the dynamic activity of protrusive structures and formation of microvilli. The video was taken with a volumetric acquisition rate of 1Hz over 300 timepoints.

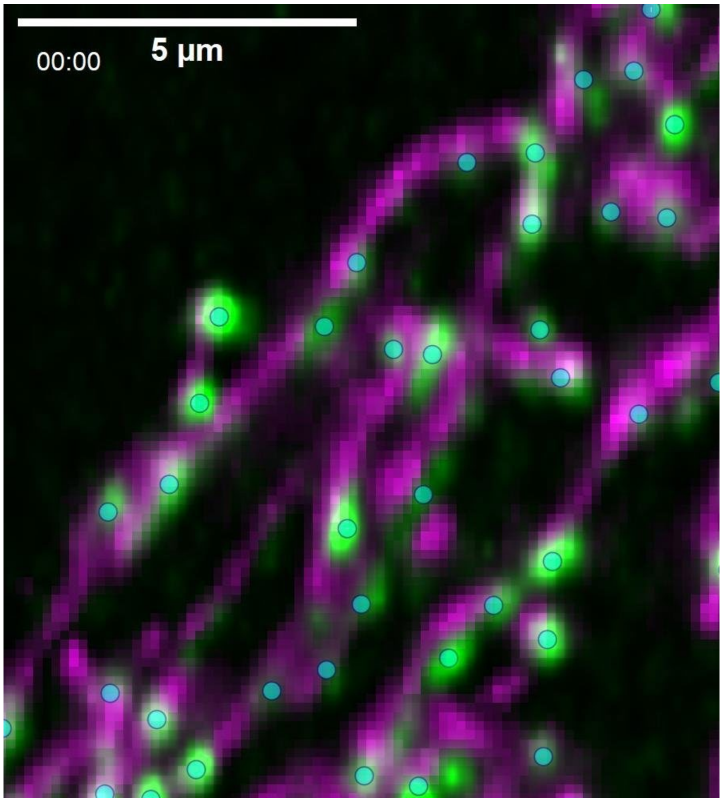


**Supplementary Video 4.** Mitochondria (magenta) and mitochondrial DNA (green) volumetrically imaged at 12s time interval. Tracking of mtDNA puncta was performed with the uTrack3D software.

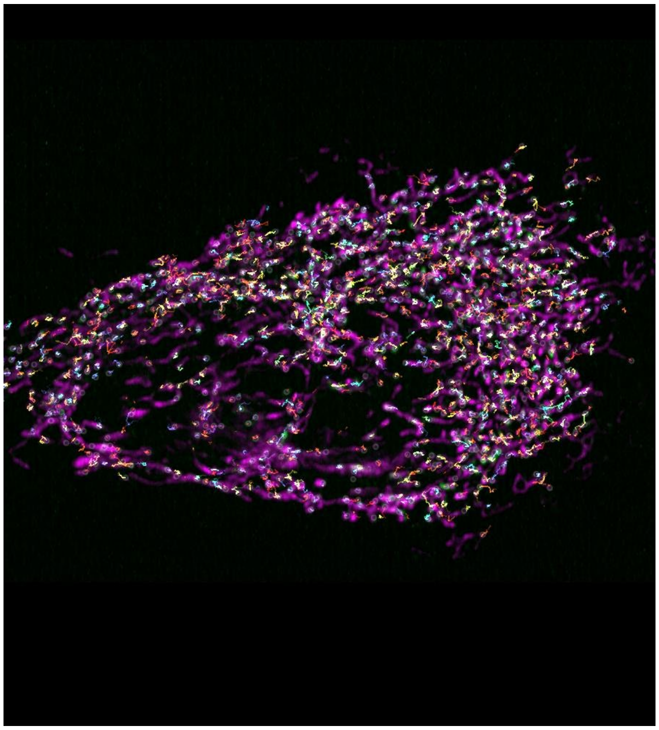

**Supplementary Video 5.** Mitochondria (magenta) and mitochondrial DNA (green) in a HeLa cell as imaged by miOPM in 5s intervals. The translocation of mtDNA along mitochondria, and apparent splitting of punctae in the event of mitochondria fission were detected and tracked using the uTrack3D software.

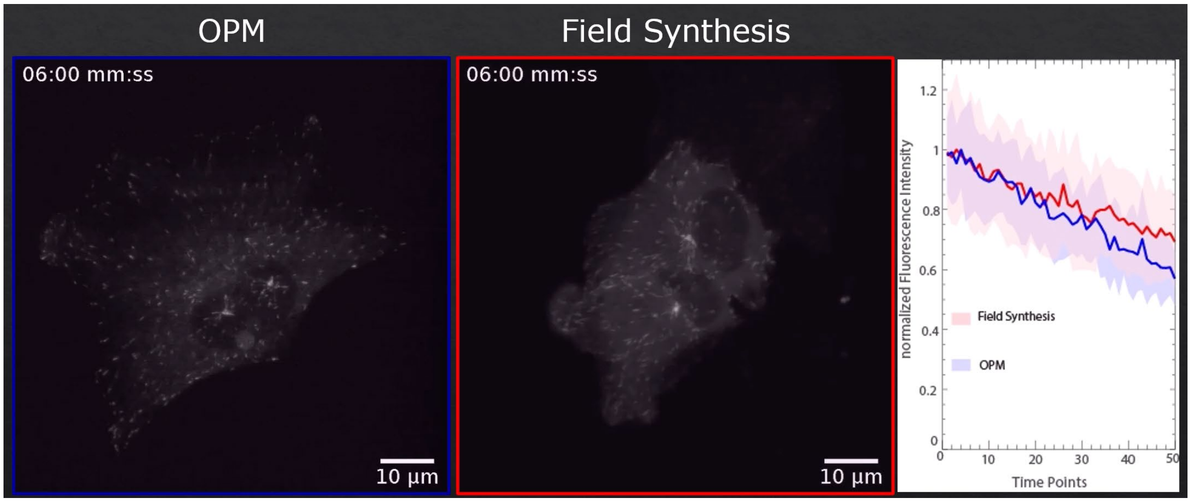


**Supplementary Video 6.** RPE cells labeled with mNeon green imaged with miOPM and a Field Synthesis microscope with a 7.5-s interval over 49 timepoints. The right side shows that bleaching rate of the two acquisitions.

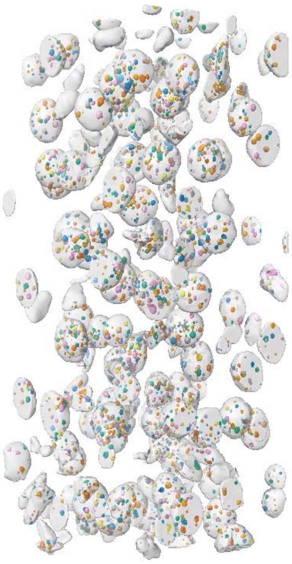

**Supplementary Video 7.** Segmentation of nuclei (labeled with sytoxGreen, rendered in gray) and nucleoli (rendered in color) in an expanded lung tissue sample as imaged by miOPM with a water immersion objective and using vertical tiling.

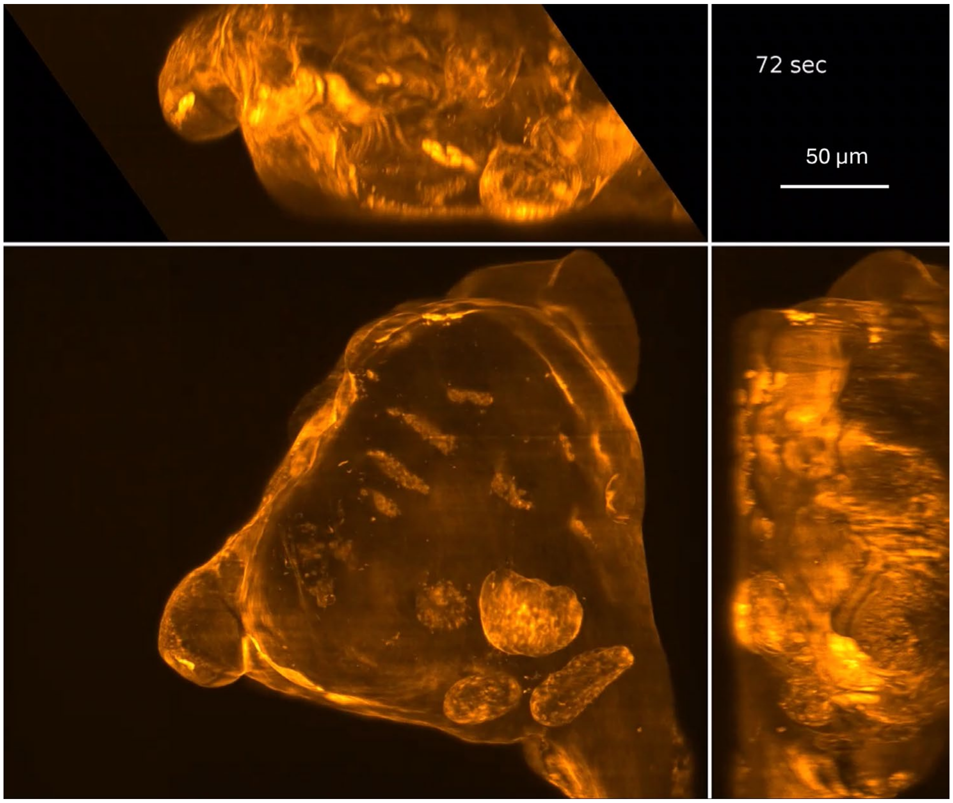


**Supplementary Video 8.** Amoeba *Chaos carolinensis* labeled with lifeact-GFP2 as imaged by miOPM with an air objective with a volumetric acquisition rate of 0.25Hz.

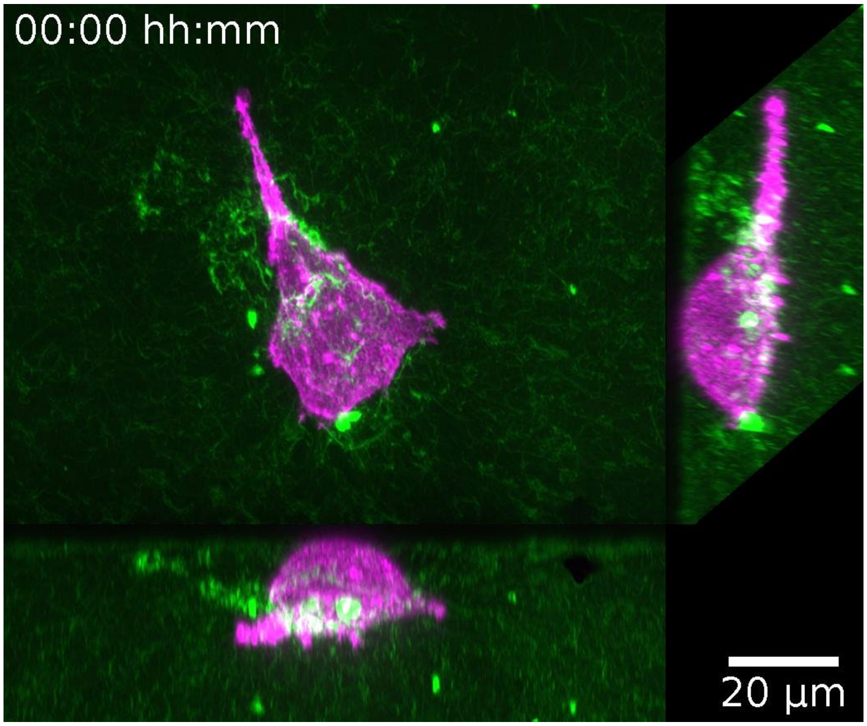


**Supplementary Video 9.** Imaging SU8686 cells labeled with Tractin-mRuby (magenta) in fluorescently labeled collagen matrix (green) in a multi-well plate over three hours.
